# *DMRscaler*: A Scale-Aware Method to Identify Regions of Differential DNA Methylation Spanning Basepair to Multi-Megabase Features

**DOI:** 10.1101/2021.02.03.428187

**Authors:** Leroy Bondhus, Angela Wei, Valerie A. Arboleda

**Affiliations:** Department of Human Genetics, David Geffen School of Medicine, UCLA, Los Angeles, CA 90095; Bioinformatics Interdepartmental PhD Program, David Geffen School of Medicine, UCLA, Los Angeles, CA 90095; Department of Pathology and Laboratory Medicine, David Geffen School of Medicine, UCLA, Los Angeles, CA 90095; Department of Computational Medicine, David Geffen School of Medicine, UCLA, Los Angeles, CA 90095; Molecular Biology Institute, UCLA, Los Angeles, CA 90095; Jonsson Comprehensive Cancer Center, UCLA, Los Angeles, CA, 90095, USA, Los Angeles, CA 90095

**Author notes:** Corresponding Author, Valerie A. Arboleda, MD PhD, 615 Charles E. Young Drive South Los Angeles, CA 90095.

**Keywords:** Rare Disease, Epigenome, DNA Methylation, Chromatin, KAT6A Syndrome, Sotos Syndrome, Weaver Syndrome

## Abstract

**Background:** Pathogenic mutations in genes that control chromatin function have been implicated in rare genetic syndromes. These chromatin modifiers exhibit extraordinary diversity in the scale of the epigenetic changes they affect, from single basepair modifications by DNMT1 to whole genome structural changes by PRM1/2. Patterns of DNA methylation are related to a diverse set of epigenetic features across this full range of epigenetic scale, making DNA methylation valuable for mapping regions of general epigenetic dysregulation. However,existing methods are unable to accurately identify regions of differential methylation across this full range of epigenetic scale directly from DNA methylation data.

**Results:** To address this, we developed DMRscaler, a novel method that uses an iterative windowing procedure to capture regions of differential DNA methylation (DMRs) ranging in size from single basepairs to whole chromosomes. We benchmarked DMRscaler against several DMR callers in simulated and natural data comparing XX and XY peripheral blood samples. DMRscaler was the only method that accurately called DMRs ranging in size from 100 bp to 1 Mb (pearson’s r = 0.96) and up to 152 Mb on the X-chromosome. We then analyzed methylation data from rare-disease cohorts that harbor chromatin modifier gene mutations in NSD1, EZH2, and KAT6A where DMRscaler identified novel DMRs spanning gene clusters involved in development.

**Conclusion:** Taken together, our results show DMRscaler is uniquely able to capture the size of DMR features across the full range of epigenetic scale and identify novel, co-regulated regions that drive epigenetic dysregulation in human disease.

## BACKGROUND

Genes that regulate chromatin structure and function are critical to coordination of complex developmental trajectories within an embryo. Mutations in these chromatin modifier genes are enriched in clinical cohorts with autism [1–4], congenital heart disease [5, 6] and global developmental delay [3, 5]. Pathogenic mutations in chromatin modifier genes can also result in specific syndromes that have both overlapping and distinct phenotypic features [7–10]. While clinical phenotypes often converge around a common set of chromatin modifier genes, the underlying molecular mechanisms driving these phenotypes are not well characterized.

Chromatin modifiers work in protein complexes to bind chromatin and shape the physical and chemical landscape of the genome, i.e. the epigenome. The regions within the genome where a particular chromatin modifier exerts its influence are critical to defining its role in development. The genomic region of affect for a chromatin modifier can be highly localized, as in methylation of individual cytosine nucleotides which modulates the binding affinity for certain transcription factors (TFs) [11–15], or it can extend across the chromatin landscape more globally, as occurs with the PRM1/2 mediated compaction of the genome during spermatogenesis [16, 17] or *Xist* in condensing the X-chromosome in cells with multiple copies of the X-chromosome [18–20]. Between the local and the global are a diversity of epigenetic features that exist at intermediate scales from tens of kilobases to many megabases. These include features such as polycomb repressive domains (PRDs) [21–23] and topologically associated domains (TADs) [24] and co-regulated gene clusters. These intermediate-sized features coordinate higher order patterning events throughout the genome in development, such as PRD regulation of *Hox* segmentation patterning [25], or organization of *Olfactory Receptor* gene clusters into TADs [26] with interdependent epigenetic regulation of the member *Olfactory Receptor* genes [27, 28]. A comprehensive understanding of chromatin modifiers requires understanding the scale of their effect on the epigenetic landscape.

While the direction of causality is still an open question for the interaction between many epigenetic features, changes in DNA methylation (DNAme) are associated with changes in other epigenetic features across the range of epigenetic scale. DNAme is the covalent addition of a methyl group to a single cytosine nucleotide usually in the context of a CpG dinucleotide [29]. While DNAme directly alters the binding affinity for a set of DNA binding proteins [11–15], it is also associated with higher order epigenetic features. At promoters and enhancers DNAme tends to be inversely correlated with gene activity [30, 31]. Over the tens to hundreds of kilobases of PRDs, DNA methylation is depleted by the antagonistic action of the polycomb repressive complex [32, 33], and as a result changes in polycomb activity over PRDs are often associated with differential methylation [33]. Megabase scale domains of active and inactive chromatin can be reliably predicted from DNAme patterns [34], and in colon cancer, changes to DNAme have been reported to overlap with these megabase-sized inactive domains [35].

Phenotypic variability and genetic heterogeneity can make the diagnosis of rare syndromes challenging. Even more challenging is the interpretation of the clinical significance of rare genetic variants identified in whole genome sequencing studies, leaving these genetic variants to be annotated as variants of unknown significance (VUSs). One method to distinguish between pathogenic and benign variants is to identify common patterns of differential DNAme from patients with known pathogenic mutations in the same gene [9, 10, 36]. The consistency of DNAme changes in these diseases suggests that common epigenetic marks are associated with specific disease phenotypes. However, directly linking observed DNAme change to the epigenetic mechanisms contributing to disease remains an open challenge.

Despite the known diversity in scale of differential DNA methylation features, no existing methods are designed to identify regions of differential methylation (DMRs) across the full range of scale from genome-wide methylation data. Instead, existing methods are designed to identify DMRs on the scale of single genes or enhancers, which provides important but incomplete information towards understanding the full epigenetic architecture. This leaves a gap in using DNA methylation to understand the dynamics of co-regulated genes and regions in a broader epigenetic context.

Here we describe a method, *DMRscaler*, that accurately identifies regions of differential methylation that can span several basepairs up to those existing at much larger scales spanning many megabases of sequence across the global DNA methylation landscape. We demonstrate the dynamic range of our differential methylation caller by simulating DMRs varying in size from 100 bp to 1 Mb and testing its performance relative to existing methods. Additionally, we use real methylation data to test for sex differences in DNA methylation where *DMRscaler*, at its highest level calls the X-chromosome as a single differentially methylated feature while still calling small, gene-level DMRs on the autosomes. Finally, we show that pathogenic mutations in chromatin modifier genes are associated with differential methylation of large and highly conserved gene-clusters such as the *HOX* and *PCDH* gene clusters. By bridging the local and the global, *DMRscaler* can provide a broadened view of differential DNA methylation structure.

## IMPLEMENTATION

To develop DMRscaler, we explored multiple pathways with the idea that we wanted to prioritize use of our method in 1) rare disease cohorts of smaller sample size and 2) be able to more readily identify the size of a differentially regulated region. In this data, DNA methylation is measured as the proportion of cytosines methylated at a given CpG site in the genome across all cells in a sample. This proportion is the β (beta) value of that site, with β=0 being completely unmethylated and β=1 being completely methylated. The distribution of β values for all CpGs across the genome follows a bimodal distribution (Additional File 1: Figure S1). Given this non-normal distribution of β values, in order to be robust in experiments with modest sample sizes (<15 samples per group, Additional File 1: Figure S2), we chose to run *DMRscaler* with the nonparametric Wilcoxon test to assign individual level CpG significance for whether the methylation distributions of the two groups differ at that CpG site. To account for multiple testing, permutation testing is used to estimate the false discovery rate (FDR) for each level of significance, and this information is used by the window scoring function (Eq. 1, described below).

To identify regions that are characterized by differentially methylated CpGs (i.e differentially methylated regions or DMRs), we used a sliding window scheme (Figure 1A,B). The window was defined by a count of adjacent CpGs rather than by the span of the genomic region. The use of a count of adjacent CpGs for window definition makes *DMRscaler* agnostic to CpG density. This allows *DMRscaler* to scan regions with low CpG coverage, such as heterochromatin, to provide suggestive evidence of differential methylation that might be missed using a distance parameter between CpGs.

**Figure 1:**
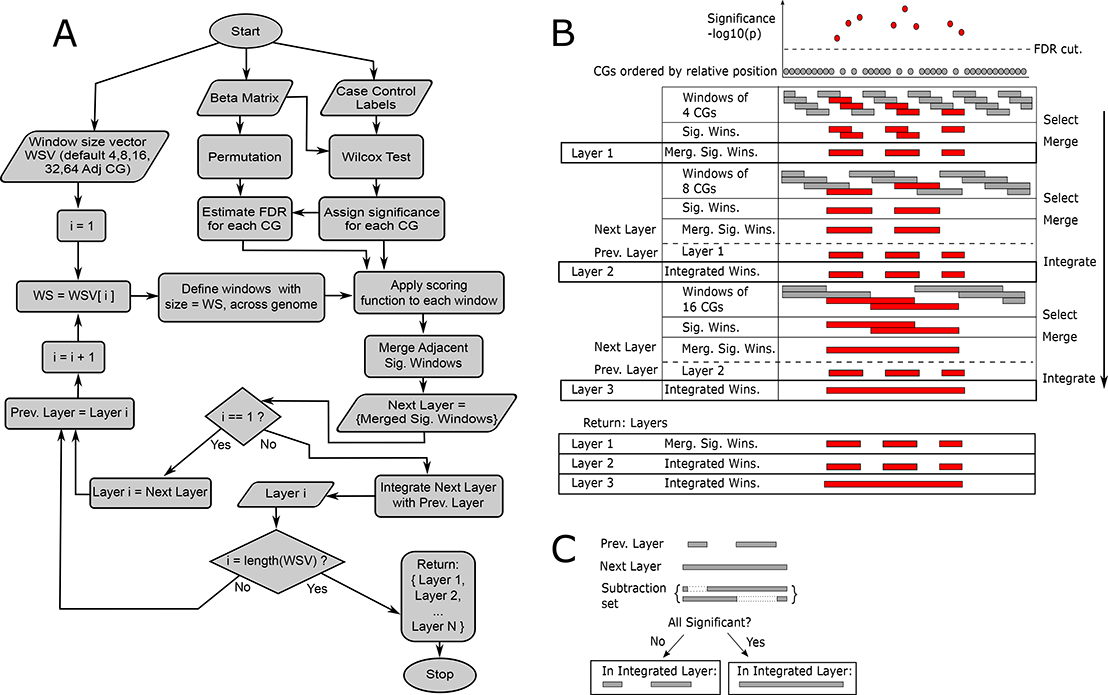
Outline of *DMRscaler* method. **(A)** Flowchart of decision tree for *DMRscaler*. Starts with Beta matrix, with individual CpG as rows and samples as columns and Beta values corresponding to methylation proportion, case-control labels, and a vector of window sizes in increasing order. Wilcoxon test and permutations are used to assign significance to CpGs as well as estimate false discovery rate FDR. The window scoring function is used with permutation to rank and assign significance to windows. Adjacent significant windows are merged forming the *Next_Layer*. For the first iteration, the returned *Layer_1* is set to this *Next_Layer*, for subsequent iterations the returned *Layer_i* is set to the result of integration of *Next_Layer* with *Prev_Layer*. Integration of layers is described in 1C. *Prev_Layer* is updated to *Layer_i* before proceeding to iteration i+1. After the largest window size layer is generated, a list is returned of the results from each iteration of the algorithm. **(B)** Graphical description of algorithm. At the top shows representation of CpGs ordered by position and associated with a significance value. Windows are laid over the ordered CpGs and selected if the window score is significant. Adjacent windows are then merged. If a *Prev_layer* has been assigned, then integration occurs. **(C)** Integration procedure. For each *Next_Layer* DMR, all overlapping *Prev_Layer* DMRs are identified. A subtraction set is generated by individually subtracting each overlapping *Prev_Layer* DMR from the *Next_Layer* DMR. Subtraction involves removing overlapping CpGs from the *Next_Layer*. If all elements of the subtraction set are significant when rescored with the window scoring function, then the *Prev_Layer* and *Next_Layer* regions are merged in the *Integrated_Layer*, otherwise the *Prev_Layer* DMRs are used in the *Integrated_Layer*. This procedure ensures that the broader *Next_Layer* DMRs are only included if no single *Prev_Layer* DMR was responsible for the significance of the region identified in the *Next_Layer*.

Once a window is defined as containing a fixed number of CpGs, the scoring function

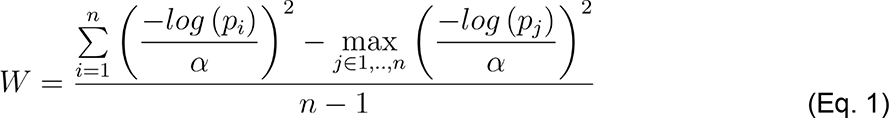

is applied to classify windows as potential DMRs. In the window scoring function, is the window score, *W* is the number of CpGs in the window, *n* is the p-value from the Wilcoxon test, and αis the significance value that corresponds to the desired FDR. Larger values of *W* suggest a window enriched in differentially methylated CpGs. Any CpGs that fall below the desired FDR significance value, will tend to weigh the overall score below 1, while those above this value, indicating evidence for differential methylation status of the window, will tend to inflate the value of . The maximum scoring CpG within a window is subtracted out so that a DMR cannot be defined by a single differentially methylated CpG. Identification of individual CpGs that are differentially methylated can be done directly from the statistical test that generated the p-value. Additionally, subtracting out the most significant CpG from the window definition is analogous to the procedure for integrating the results of DMRs called using increasing window sizes detailed below, suggesting a general integration strategy.

After windows are scored, a permutation scheme is applied to estimate the significance of the score of each window as the expected frequency at which a random grouping of CpGs would have resulted in the given window score or greater. This permutation uses the p-value assigned to individual CpGs in the true case v control partition. Windows exceeding a preselected significance value (significant windows) are merged with adjacent and overlapping significant windows and retained as candidate DMRs. The p-value cutoff for windows was set empirically to p-value=1E-4. At this value the expected false positive rate was FDR < 0.1 for each window layer for each analysis based on the expected number of windows identified as significant and the returned number of significant windows (Additional File 1: Table S1).

In order to define DMRs that can span whole chromosomes (see Results) as well as those that are much narrower, such as those limited to individual promoters, within the same analysis, we iteratively identify candidate DMRs using windows of increasing size (defined as layers) and then integrate the results across these layers as a means to identify the scale of the DMR (Figure 1B,C). The smallest window size is used first to build *layer_1_* in the output. As a note, the term ’layer’ is used to describe the resulting set of DMRs constructed with a given window size parameter to suggest the relation of the results of each iteration of the algorithm. The construction of each successive layer either expands, adds, or retains DMRs from the *previous layer*, and so there is a hierarchical relation between DMRs across layers. DMRs in lower layers will always map to some DMR in upper layers, ensuring that *layer_i_* will always be a subset of *layer_i+1_*. To define the *next layer*, the following steps are repeated for windows of the next largest size: overlaying windows, identifying windows significantly enriched in differentially methylated CpGs, and merging significant windows (Figure 1A,B). From the second layer onward, an additional step to integrate the results from the *previous layer* is performed. This is achieved by subtracting each *previous layer* DMR from any overlapping DMR in the *next layer*, and retesting the reduced DMR in the *next layer* for significance (Figure 1C). If a DMR in the *next layer* remains significant following the subtraction of each overlapping *previous layer* DMR individually, then the *next layer* DMR is retained in the new *integrated layer*. Additionally, any *previous layer* DMRs that did not overlap a DMR in the *next layer* are added to the *integrated layer*. In this way, at each iteration of the algorithm DMRs are either added, expanded, or consolidated from the *previous layer* to the *integrated layer* but never lost. As the algorithm proceeds, the *previous layer* is updated to the most recent *integrated layer* before the next step of integration is done with the *next layer*. Through iteratively calling DMRs using windows of increasing size and integrating the results, *DMRscaler* is able to identify DMRs that vary dramatically in terms of scale.

## METHODS

### Cell culture

For *KAT6A* data, fibroblast cell lines were derived from skin punch biopsies performed on the proband and one or both unaffected parents. This project was approved by the UCLA Institutional Review Board #11-001087. All individual level-data was de-identified prior to analysis. Fibroblast cell culture lines were created through the UCLA Pathology Research Portal and fibroblast cell lines were established and grown in DMEM (Gibco™), 10% FBS (Heat-inactivated Fetal Bovine Serum, Thermofisher), 1% Non-essential Amino Acid (Gibco™) and 1% PenStrep at 37℃ in 5% CO_2_ incubators. Cell lines were tested for mycoplasma at least on a monthly basis.

### DNA methylation studies

For KAT6A methylation studies, DNA was extracted from patient-derived fibroblast cell lines. The specific mutation for each line is given in Additional File 1: Table S2. DNA samples were bi-sulfite converted and run on the Illumina MethylationEPIC Array (850k EPIC array) as previously described [37] at the UCLA Neuroscience Genomics Core to generate idat files. QC on the resulting idat files was done using the MINFI package, and probes overlapping SNPs were removed [38]. After QC, 852,671 of 865,919 measured CpGs remiained, after removal of sex chromosome CpGs, 832,159 measured CpGs remained. Preprocessing and normalization of individual probes was done using background correction [39] and functional normalization [40].

### Data sources

Publically available datasets of peripheral blood methylation data for control, Weaver Syndrome and Sotos Syndrome patients were downloaded from the Gene Expression Omnibus (GEO) resource [41, 42] with accession number GSE74432 [10].

### Simulation

To demonstrate how *DMRscaler* distinguishes itself from other methods, we simulated differentially methylated regions (DMRs) ranging in size across several orders of magnitude (Figure 2A).

**Figure 2:**
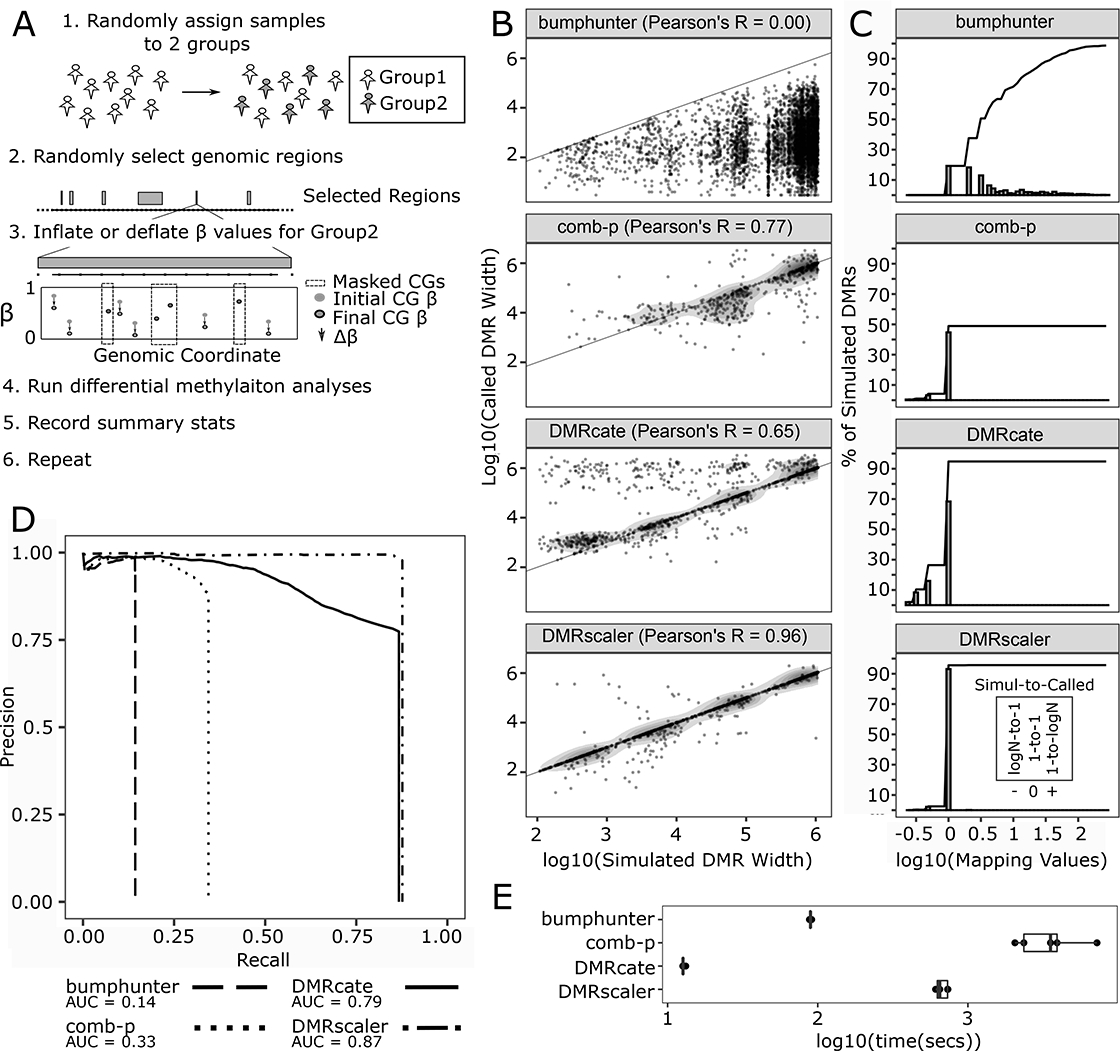
Simulation of DMRs ranging in size between 1kb to 1Mb for comparison of methods. (A) Graphical description of simulation design. First, samples are randomly assigned to one of two groups. Second, non-overlapping regions of the genome are randomly selected to be DMRs. Third, over selected DMRs one group has the β value of non-masked CpGs inflated or deflated by Δβ. Next all differential methylation methods are run and relevant summary statistics are recorded. This procedure is repeated a number of times to generate additional data points. (B) Simulated DMR Widths v Called DMR Widths plotted on log10 scale. Pairs are formed between simulated and called DMRs if there is any overlap between the two. (C) Mapping Values plots. The mapping value is calculated for each simulated DMR and is either the inverse of the number of simulated DMRs sharing an overlapping called DMR or else it is the number of called DMRs overlapping the given simulated DMR, whichever is more extreme. Log values > 0 imply multiple DMRs called per simulated DMR. Value < 0 imply multiple simulated DMRs overlap single called DMR. Value = 0 implies one DMR called per DMR simulated. The plotted line indicates the cumulative proportion of simulated DMRs up to the given mapping value. (D) Feature level Precision-Recall Curves for each method, see methods for details on calculation. (E) Time for each method to run on the simulated dataset across 5 runs.

DNA methylation measured on the Infinium HumanMethylation450 BeadChip (450K array) from whole blood for 53 controls from GEO (GSE74432) was used as a foundation for the simulation [10]. Real data was used as the foundation in order to capture the natural biological and technical variability present in DNA methylation array data. QC on the resulting idat files was done using the MINFI package, and probes overlapping SNPs were removed [38]. After QC, 468,162 of 485,512 measured CpGs remained, and after removal of the sex chromosomes 456,514 measured CpGs remained. Preprocessing and normalization of individual probes was done using background correction [39] and functional normalization [40].

Regions for artificially introducing DMRs were selected at random across the genome but subject to the following constraints. DMRs specified as 0.1-1 kb in size were required to have at least 3 CpGs represented on the 450K array (CpGs), those 1-10 kb in size were required to have at least 6 CpGs, those 10-100 kb in size were required to have at least 9 CpGs, those 0.1-1 Mb in size were required to have at least 12 CpGs. Additionally, to avoid miscounting, DMRs were introduced such that they were spaced at least 10 CpGs apart from any other introduced DMR.

All 450k array samples used were from control whole blood, so for each run of the simulation samples were pulled at random into one of two groups, Group1 and Group2. Each group consisted of 8 samples drawn without replacement from the pool of 53 samples.

Before artificially introducing the DMRs to the real data matrix, 50% of CpGs within each DMR, excluding the first and last CpG, were randomly masked and kept at their original values. This was done to model the variability of methylation state of neighboring CpGs in real data. Then, the mean β value of CpGs within each DMR was measured for Group1 and Group2. The β values of the group with the greater mean β value would have all non-masked CpGs inflated by 0.2 to model a modest effect DMR. If this resulted in any samples having a β value greater than 1, the β values for that CpG were divided by the max β value for that CpG to bring values back to the range of 0-1. Other values of inflation for β were also tested without notable changes in the results (Additional File 1: Table S3).

Following the introduction of artificial DMRs into the dataset, *DMRscaler*, *bumphunter* [43]*, comb-p* [44] and *DMRcate* [45] were run on the dataset and the results tabulated. *DMRscaler* was tested using a window_size_vector of 4, 8, 16, 32, 64 Adj CpGs, FDR threshold of 0.1, window_p_cut of 1e-4 corresponding to empirically determined FDR of <0.1 from real data analysis (Additional File 1: >Table S1). *Bumphunter* was tested with MaxGap = 1e6 with loess smoothing enabled. *Comb-p* was tested with dist = 1e6, step = 5000, seed = 1e-3, region-filter-p = 0.1 (Additional File 1: Figure S3). *DMRcate* was tested with lambda = 1e6, C=2000. Parameter sets for methods were chosen to facilitate identification of larger DMRs for output more comparable to *DMRscaler*.

The area under the precision-recall curve (AUCPR) (Figure 2D) was measured at feature level following the framing of the problem for measuring precision and recall for time series proposed by Tatbul and colleagues, which is generally appropriate for other forms of range data when identification of individual features is of interest [46]. Each simulated DMR was considered as a true feature with true positive (TP) and false negative (FN) attributes. Each called DMR was considered a called feature with TP and false positive (FP) attributes. Called DMRs were ordered by their score for *DMRscaler* and p-value for *bumphunter, comb-p,* and *DMRcate*. The PR curve was generated by measuring precision (P) and recall (R) with stepwise inclusion of the next highest scoring or most significant called DMR. At the n-th step precision and recall were measured as

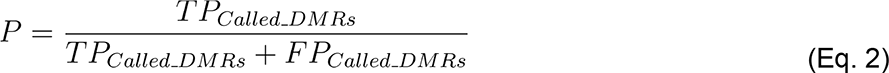

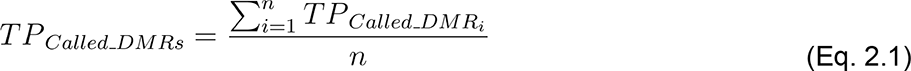

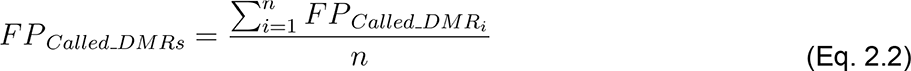

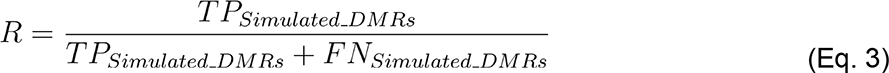

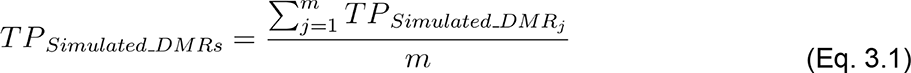

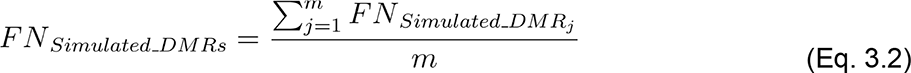

where *TP _called_DMRi_* is the proportion of the i-th called DMR overlapping simulated DMR regions, *FP_Called_DMRi_* is the proportion of the i-th called DMR not overlapping a simulated DMR region, *TP_Simulated_DMRi_* is the proportion of the j-th simulated DMR overlapping any of the 1 to n-th Called DMRs, *FN_Simulated_DMRi_* is the proportion of the j-th simulated DMR overlapping any of the 1 to n-th Called DMRs, is the number of most significant called DMRs used at the n-th step, and is the total number of simulated DMRs. The feature level measure of precision and recall gives equal weight to each simulated DMR so that large simulated DMRs do not dominate the signal.

### Rare Disease Data Analyses

For each real data analysis, *DMRscaler*, *bumphunter, comb-p*, and *DMRcate* were used to call DMRs. *DMRscaler* was tested using a window_size_vector of 4, 8, 16, 32, 64 Adj CpGs, FDR threshold of 0.1, window_p_cut of 1e-4 corresponding to an empirically determined FDR of < 0.1 (Additional File 1:Table S1). *DMRcate* was tested with default parameters, as well as with lambda = 1e6, C=2000 to capture larger DMRs for output more comparable to *DMRscaler*. *Bumphunter* was tested with default parameters, as well as with MaxGap = 1e6 with loess smoothing enabled.

For the sex analysis, DNA methylation measured on the Infinium HumanMethylation450 BeadChip (450K array) from whole blood for 53 controls from GEO (GSE74432) was used [10]. QC on the raw idat files was done using the MINFI package, and probes overlapping SNPs were removed [38]. After QC, 468,162 of 485,512 measured CpGs remained. Preprocessing and normalization of individual probes was done using background correction [39] and functional normalization [40]. A subsample of 8 control males and 8 control females from this dataset was used for analysis to be more comparable to a typical rare disease study design. Raw output from each method is provided in Additional File 2: Table S4.

For KAT6A analysis, DNA methylation was measured on the Illumina MethylationEPIC Array (850k EPIC array) with 8 cases and 12 controls. QC on the resulting idat files was done using the MINFI package, and probes overlapping SNPs were removed [38]. After QC, 852,671 of 865,919 measured CpGs remained, after removal of sex chromosome CpGs 832,159 measured CpGs remained. Preprocessing and normalization of individual probes was done using background correction [39] and functional normalization [40]. Raw output from each method is provided in Additional File 3: Table S5.

For Weaver analysis, DNA methylation measured on the Infinium HumanMethylation450 BeadChip (450K array) from whole blood for 8 patients with *EZH2* mutations and 53 controls from GEO (GSE74432) was used [10]. This data comes from a study that found an epigenetic signature specific to Sotos syndrome from *NSD1* mutations using Weaver syndrome samples as a negative control for their classifier [10]. More recently, this data has been used to identify an epigenetic signature specific to Weaver syndrome [9]. QC on the raw idat files was done using the MINFI package, and probes overlapping SNPs were removed [38]. After QC, 468,162 of 485,512 measured CpGs remained, and after removal of the sex chromosomes 456,514 measured CpGs remiained.. Preprocessing and normalization of individual probes was done using background correction [39] and functional normalization [40]. Raw output from each method is provided in Additional File 4: Table S6.

For Sotos syndrome analysis, DNA methylation measured on the Infinium HumanMethylation450 BeadChip (450K array) from whole blood for 38 patients with *NSD1* mutations and 53 controls from GEO (GSE74432) was used [10]. QC on the raw idat files was done using the MINFI package, and probes overlapping SNPs were removed [38]. This comes from the same study as the Weaver syndrome data [10]. After QC, 468,162 of 485,512 measured CpGs remained, and after removal of the sex chromosomes 456,514 measured CpGs remiained. Preprocessing and normalization of individual probes was done using background correction [39] and functional normalization [40]. Raw output from each method is provided in Additional File 5: Table S7.

### Syndrome DMR Overlap Analysis

To test for overlapping regions of differential methylation between KAT6A, Sotos, and Weaver syndrome, the number of measured CpGs considered for DMR detection was downsampled to include only those CpGs measured on both the Infinium HumanMethylation450 BeadChip (450K array) and the Illumina MethylationEPIC Array (850k EPIC array). This left 425,733 measured CpGs for calling DMRs.

Overlaps between DMRs were counted between syndromes as was the overlap of gene sets. Gene set overlaps were considered separately to identify genes that may be commonly differentially methylated but identified by non-overlapping regions of the gene, something the direct DMR overlap measure would miss. Raw output of region level and gene level overlaps are provided in Additional File 6: Table S8.

To test whether CpGs identified as belonging to DMRs are enriched between syndromes, that is whether membership of a CpG to a DMR in one syndrome makes it more or less likely to also belong to a DMR in another syndrome, we computed the odds ratios (OR). The OR was calculated by forming a 2x2 contingency table with counts of CpGs belonging to DMRs in both syndromes, CpGs belonging to one and not the other, and CpGs belonging to DMRs in neither. The raw counts used in the 2x2 contingency tables are given in Additional File 1: Table S9. Odds ratios for all pairs of syndromes are given in Additional File 1: Table S10.

## RESULTS

### DMRscaler Overview

Our goal in developing *DMRscaler* was to have a method capable of accurately identifying regions that demonstrate differential methylation across the full range of epigenetic scale, from small-promoter to whole-chromosome scale features. The major bottleneck to this goal is that regions of differential methylation show significant variability in methylation state between neighboring CpGs. For example, nearly 20% of neighboring CpGs between 0.5 - 1.0 kb away have a difference in the proportion of methylation greater than 50% (Additional File 1: Figure S4). When trying to identify DMRs that may span larger genomic regions, such as gene clusters, this variability makes the trivial method of taking contiguous blocks of significant CpGs as the DMRs ineffective. One approach to resolve this issue of high variability is to smooth differential methylation sites based on significance across adjacent CpGs or over some specified genomic interval. However, the smoothing approach is sensitive to the choice of bandwidth parameter used for the smoothing window. Windows that are too small will fail to connect features over larger gaps, windows that are too large will result in excessively broad DMRs. Smoothing alone is therefore inappropriate when features are expected to vary dramatically in terms of scale. To capture potentially noisy features that may vary in size by several orders of magnitude, from the basepair to multi-megabase scale, we need a method that is both robust to noise and that can accurately determine the feature’s size.

To address these limitations in determining the size of a DMR, *DMRscaler* uses an iterative sliding window over the genome (Figure 1A,B), represented as a partially-ordered set of measured CpGs, and implements an integration step between each iteration of the sliding window (Figure 1C). The windows at each step identify the set of regions that are enriched in CpGs with significantly different methylation values between cases and controls. By binning CpGs into windows and testing these windows for enrichment in significant CpGs (Eq. 1), the algorithm is robust to noise caused by variability in methylation of neighboring CpGs. To address the bias in feature size introduced by preselecting a window size parameter, *DMRscaler* calls significant windows iteratively with a variable increasing size parameter and integrates the result of each iteration with the results from the previous iterations. The integration step (Figure 1C) is used between the previous (lower) layer, built from smaller windows, and the current (upper) layer to determine which features in the upper layer are already adequately represented by lower layer features and which upper layer features capture a statistically significant association missed by the lower layer features. If an upper layer feature captures a statistically significant association missed in the lower layer then that upper layer feature is retained and resolved with any overlapping lower layer features, otherwise the overlapping lower layer representation is carried through unmodified. For a more detailed description, see implementation.

*DMRscaler* provides a solution to the problem of identifying DMR features across the full range of epigenetic feature sizes, whether at the basepair level or across entire chromosomes. The integration of results across iterations of the windowing procedure *DMRscaler* implements is a novel mechanism for defining DMRs that could be generalized to other epigenetic features or one dimensional data where discontinuity in components defining a feature of interest is expected.

### Comparison of DMRscaler with existing methods

We next benchmarked *DMRscaler* to three commonly used methods in identification of differentially methylated regions: *bumphunter* [43], *comb-p* [44], and *DMRcate* [45] (Table 1). One significant difference is that our approach uses Wilcoxon’s Test to estimate significance of individual CpGs while *bumphunter* and *DMRcate* use a t-test. *Comb-p* does not estimate individual CpG significance but instead takes this as an input value. We observed that when running with a small sample size (n=8 per group) there is poor correlation between the significance of differential methylation determined by the Wilcoxon and t-test (Additional File 1: Figure S2). Since one of our goals was to develop a method that could detect DMRs in studies that compare rare disease datasets, we sought to implement a test that worked with non-normal data distributions (Additional File 1: Figure S2). While the t-test is appropriate when the sample size is sufficiently large (n > 30) or the sampling distribution is approximately normal, differential methylation analysis in small samples breaks these assumptions and therefore in our use-case the Wilcoxon test is more appropriate.

**Table 1:** Comparison of Differential Methylation Methods

|  | *DMRscaler* | *bumphunter* | *comb-p* | *DMRcate* |
| --- | --- | --- | --- | --- |
| Individual CpG significance | Mann-Whitney Test | T-Test | NA (takes p-value as input) | T-Test, using M transformed Beta values |
| DMRs definition | Iterative enrichment testing for significant CpGs within window. | Consecutive CpGs above significance threshold | Groups significant CGs if within window or window interval | Gaussian smoothed regions above significance threshold |
| DMR significance | Permutation Test | Stouffer's Method | Stouffer-Liptak | Permutation Test |
| Parameters controlling size (default) | window_size (4,8,16,32,64 adj CpGs) | maxGap (500 bp) | dist, step (no default) | lambda (1 kb), C (2) |

The second difference between the differential methylation callers compared here (Table 1) lie in their modeling to identify differentially methylated regions. Briefly, *bumphunter* uses a linear regression model to identify CpG sites that are differentially methylated between case and control conditions. Then to detect DMRs, *bumphunter* identifies stretches of adjacent CpGs that are above a specified significance threshold. However, the methylation landscape of adjacent CpGs is complex, with CpGs with high, intermediate and low β values mixed together making definition of large and contiguous regions of differential methylation challenging (Additional File 1: Figure S4, S5). *Comb-p* uses the Stouffer-Liptak method for p-value correction and then groups significant CpGs within a window or window interval defined by the *dist* and *step* parameters. *DMRcate* is similar to *bumphunter* in that it also implements linear modeling (Table 1). It differs from *bumphunter* in that rather than implementing a strict adjacency requirement, *DMRcate* uses a Guassian smoothing function on M transformed β values to identify DMRs in genome-wide data. This provides the user with control of a bandwidth parameter, lambda, and control parameter, C, that can be used to identify larger regions of differential methylation. However, the behavior of *DMRcate* at larger bandwidth is poorly defined and the size of DMRs returned tends to be sensitive to parameter choice. For an in-depth review of methods see [47].

The design of the *DMRscaler* method has several unique features that allow it to more accurately identify larger co-regulated regions. First, it incorporates the intrinsic variability in methylation distribution across the genome by binning adjacent CpGs into windows before assigning significance. Second, *DMRscaler* integrates the results from layers of windows defined with a series of window sizes. To accommodate the study of rare-disease cohorts where sample availability is often limited, our study uses Wilcoxon’s test for p-value significance which is appropriate for small datasets with non-normal distributions, however other methods of generating p-values could be used, for instance to model the effect of covariates. Together, these features allow for the robust detection of differentially methylated regions across a large dynamic range, spanning basepair to megabase resolution and allow for detection of novel regions that are differentially methylated in rare disease cohorts and between sexes.

### *DMRscaler* accurately captures the scale of epigenetic features from basepair (bp) to megabase (Mb) size in simulated methylation data

To benchmark our method against existing methods, we compared *DMRscaler* to *bumphunter, comb-p,* and *DMRcate* on metrics including: the correlation between simulated DMRs and DMRs called by each method across a wide rage of simulated DMR sizes, the mapping value or the degree to which each method was able to represent individual simulated DMRs as single unified features, the precision and recall of simulated DMRs, and the run time of each method.

We first simulated DMRs in methylation data from control blood samples (GSE74432) [10] ranging in size from 100 bp to 1 Mb (Figure 2A, see method for details). In our simulation, we modelled noise within DMRs to represent the situation observed in real data where neighboring CpGs often have distinct methylation states.

*DMRscaler* was able to accurately call the size of the simulated DMRs (pearson’s r = 0.96) relative to *bumphunter* (pearson’s r = 0.00), *comb-p* (pearson’s r = 0.77), and *DMRcate* (pearson’s r = 0.65) (Figure 2B). *DMRscaler* preserves a strong 1-to-1 relation between simulated and called DMRs, with 93% of simulated DMRs accurately called by *DMRscaler* with a 1-to-1 relation, compared with 19% for *bumphunter*, 45% for *comb-p*, and 68% for *DMRcate* (Figure 2C).

To measure performance of our differential methylation caller, we calculate the AUCPR for each test. AUCPR combines a measure of precision of features called (ratio of true feature called to all features called) and recall (ratio of true features called to total number of true features) into a single value, with AUCPR = 0 representing no classification and AUCPR = 1 representing perfect classification of all features with no false positives. In our simulation, *DMRscaler* had an AUCPR of 0.87, *bumphunter* had an AUCPR value of 0.14, *comb-p* had an AUCPR of 0.33, and *DMRcate* had an AUCPR of 0.79 (Figure 2D, see methods for details on AUCPR calculation). The low AUCPR of *bumphunter* is consistent with the low, slightly negative correlation observed between simulated and called DMR regions (Figure 2B). This weak correlation is due to the fact that *bumphunter* has a strict requirement that significant differentially methylated CpGs are adjacent in order to belong to a common DMR and therefore breaks up simulated DMR features into many smaller features. The low AUCPR of *comb-p* is the result of a low recall rate of features that are much smaller than the size set by the *dist* and *step* parameters. Setting lower values for the *dist* parameters increases the ability to detect smaller DMR features but at the expense of detecting larger DMR features (Additional File 1: Figure S3), and at very large values for *dist* the run time becomes prohibitive especially as smaller step values are used (Additional File 1: Figure S3). *DMRcate* had a reasonably high AUCPR, however there is a bias in the size of DMRs called based on the choice of the bandwidth parameter lambda, and the control parameter C. Specifically, there is an excess of false calls of DMRs around 1 Mb and 1 kb (Figure 2B and S6) which is related to the choice of bandwidth parameter λ (set to 1 Mb) and the scaling parameter C (ratio of λ/C set to 500) (Additional File 1: Figure S7). Our data suggests that *DMRcate* is able to identify larger DMRs but also that called DMR size is sensitive to parameter choice for the lambda and C parameters. This is supported by the shape of the precision-recall curve for *DMRcate* that shows a modest drop in precision as recall increases, suggesting *DMRcate* incurs a steeper false positive penalty compared with *DMRscaler*.

While *DMRscaler* performs well compared to other methods at the task of identifying DMRs across a wide range of scales, the method is relatively slow. On average *DMRscaler* required 25-30 min to complete a run, *bumphunter* required only a little over 1 min, and *DMRcate* only required around 10 seconds to call DMRs. *Comb-p,* which uses a sliding window mechanism similar to *DMRscaler,* required around an hour to complete each run with the given parameter set (Figure 2E).

The simulation results show that *DMRscaler*, while not optimized for speed, reconstructs the scale of DMR features more accurately than other methods across a wide range of DMR feature sizes as measured by called and simulated DMR size correlation, mapping value, and precision and recall .

### Differential methylation between sexes captures chromosome-wide and gene specific regulatory features in empiric data

To test our hypothesis in real-world DNA methylation data, we sought to determine whether our method could capture both small regions of autosomal differential methylation as well as chromosome-wide features such as X-chromosome inactivation. Therefore, our test case is the natural occurrence of X-inactivation in females, where one copy of the X-chromosome is largely inactivated by the action of the lncRNA *Xist* [18, 19]. This process of inactivation is correlated with a striking chromosome wide difference in DNA methylation between males and females on the X-chromosome (Figure 3A top) as compared to the autosomes, e.g. chromosome 1 where the size of differentially methylated regions span 166 bp to 512 bp (Figure 3A bottom, Additional File 2: Table S4). Across all chromosomes, the size of DMRs called by *DMRscaler* spans 86 bp to 152 Mbp, representing a 1,768,750- fold difference in scale detected by *DMRscaler*.

**Figure 3:**
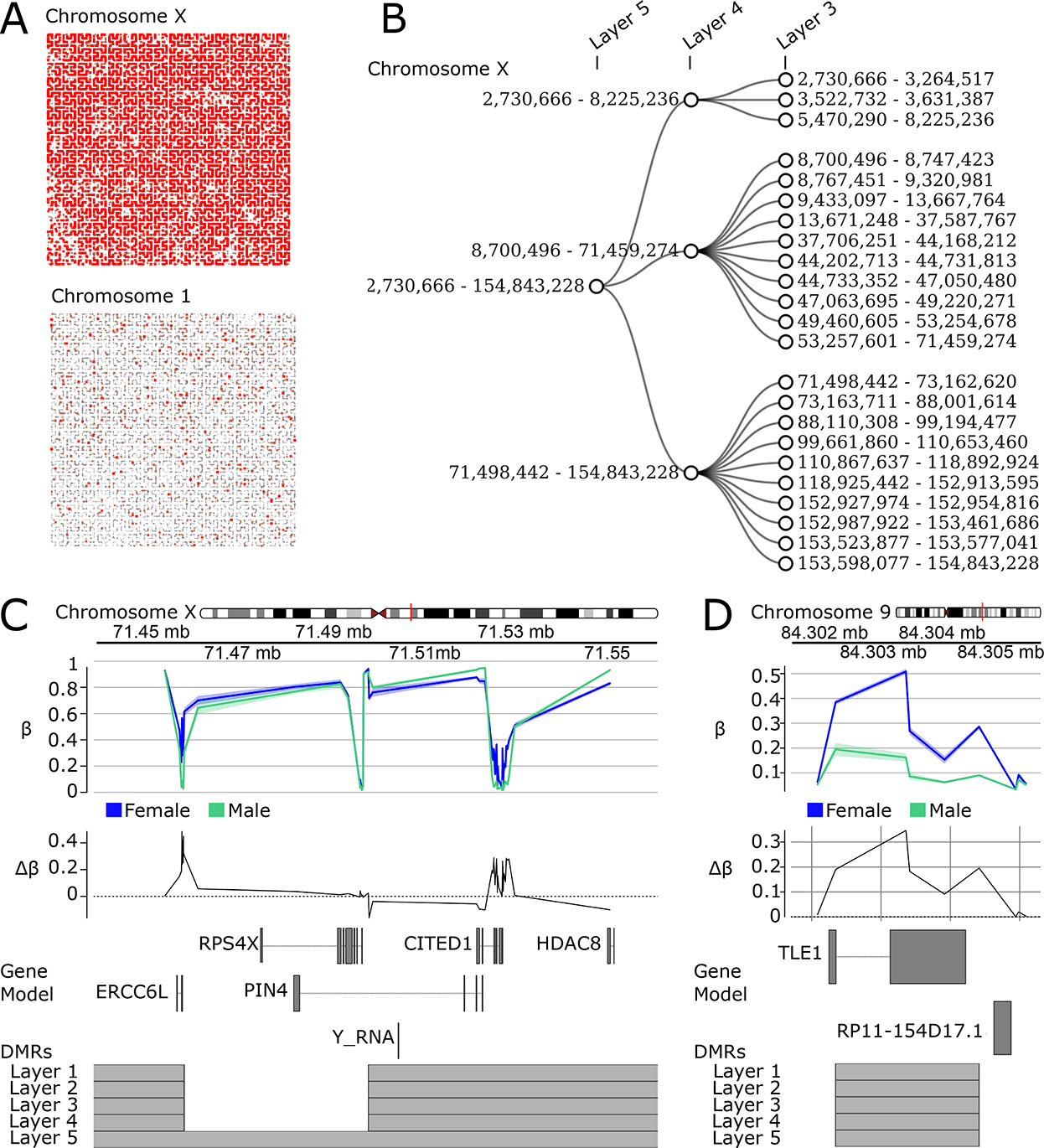
Differential Methylation Analysis Between XX and XY individuals. **(A)** Hilbert Curves of chrX and chr1. Hilbert curve is constructed by ordering CpGs by their position along the given chromosome. Red points are differentially methylated CpGs with FDR < 0.1. Point size scaled to max significance level (-log10 p-value). **(B)** Diagonal network plots showing hierarchical relation of DMRs called by *DMRscaler* in Layers 5,4,3 (equivalent to 64,32,16 Adj. CpG Layers respectively) for X-chromosome. **(C)** ChrX:71.4-71.6Mb. GVIZ track stack plot. Top track shows mean β value per group, next track shows Δβ, where Δβ = β_female_ - β_male_ . Below the gene model track is the DMR track, highlighting the regions called as a DMR at each result layer from *DMRscaler* (Layers 1, 2, 3, 4, 5 are equivalent to 4, 8, 16, 32, 64 Adj. CpG layers). (D) Chr9:84.302-84.306Mb. Tracks same as 3C.

With the visual intuition of the scale of differential methylation between sexes from Figure 3A, we next compared the result of differential methylation analysis using *DMRscaler*, *bumphunter*, *comb-p* and *DMRcate*. *DMRscaler* was the only method that consolidated the observed differential methylation into a single DMR that spanned 98% of the X-chromosome (Table 2, Figure 3B, Additional File 1: Figure S8). Even with the maxWidth parameter set to 1 Mb, *Bumphunter* reported 1,231 unique DMRs on the X-chromosome with a median width of 2.8 bp (IQR: 1 bp - 446 bp) (Additional File 1: Figure S9, Additional File 2: Table S4), likely due to a lack of mechanism for spanning non-differentially methylated CpGs. With a standard parameter set of dist = 1 kb, step = 100bp, *comb-p* reported 866 unique DMRs on the X-chromsome with a median width of 666 bp (IQR: 331 bp - 1.08 kb) (Table 2, Additional File 1: Figure S9, Additional File 2: Table S4). With a wider parameter set of dist = 1 Mb, step = 100 kb, *comb-p* called 19 unique DMRs on the X-chromosome with a median width of 3.15 Mb (IQR: 512 kb - 8.53 Mb) (Table 2, Additional File 1: Figure S9, Additional File 2: Table S4 ) . *DMRcate* with default settings reported 1,062 unique DMRs on the X-chromosome with a median width of 1.08 kb (IQR: 615 bp - 1.68 kb). When *DMRcate* was provided with a larger bandwidth parameter (lambda = 1 Mb, C = 2000) it improved in consolidating the DMRs, but still reported 19 unique DMRs (median width: 2.64 Mb, IQR: 0.82 Mb - 8.25 Mb). This improvement in consolidation of the X-chromosome was accompanied by a large increase in the average size of autosomal DMRs called (mean 173.9 kb, up from 644 bp with default parameters) and total autosomal genome covered by DMRs called (1.8%, up from 0.0098% with default parameters) (for complete distributions see Additional File 1: Figure S10, Additional File 2: Table S4).

**Table 2:** Sex Analysis DMR Summary Table

| Sex Analysis DMR Summary Table |  |  |  |  |  |  |  |  |
| --- | --- | --- | --- | --- | --- | --- | --- | --- |
| Method | Chromosome X |  |  |  | Autosomes |  |  |  |
|  | # DMRs | Mean DMR width | Median DMR width | % total width | # DMRs | Mean DMR width | Median DMR width | % total width |
| *DMRscaler* |  |  |  |  |  |  |  |  |
| Layer 1: 4 adj CpGs | 280 | 364.02 kb | 160.51 kb | 66% | 28 | 875 bp | 441 bp | 0.00085% |
| Layer 2: 8 adj CpGs | 174 | 643.80 kb | 203.00 kb | 72% | 28 | 915 bp | 441 bp | 0.00089% |
| Layer 3: 16 adj CpGs | 23 | 6.49 Mb | 2.75 Mb | 95% | 30 | 3.75 kb | 477 bp | 0.0039% |
| Layer 4: 32 adj CpGs | 3 | 50.53 Mb | 62.76 Mb | 98% | 31 | 31.08 kb | 512 bp | 0.033% |
| Layer 5: 64 adj CpGs | 1 | 152.11 Mb | 152.11 Mb | 98% | 31 | 31.08 kb | 512 bp | 0.033% |
| *bumphunter* |  |  |  |  |  |  |  |  |
| Default:<br>maxGap = 500 bp | 1452 | 269 bp | 28 bp | 0.25% | 104 | 14 bp | 1 bp | 0.000049% |
| maxGap = 1 Mb | 1231 | 2.48 kb | 135 bp | 2.0% | 34 | 34 bp | 1 bp | 0.00012% |
| *comb-p* |  |  |  |  |  |  |  |  |
| dist = 1 kb,<br>step= 100 bp | 1452 | 269 bp | 28 bp | 0.25% | 104 | 14 bp | 1 bp | 0.000049% |
| dist = 1 Mb,<br>step | 1231 | 2.48 kb | 135 bp | 2.0% | 34 | 34 bp | 1 bp | 0.00012% |
| *DMRcate* |  |  |  |  |  |  |  |  |
| Default:<br>lambda = 1kb,<br>C = 2 | 1062 | 1.31 kb | 1.08 kb | 0.90% | 439 | 644 bp | 572 bp | 0.0098% |
| Lambda = 1 Mb,<br>C = 2000 | 19 | 6.16 Mb | 2.64 Mb | 76% | 291 | 173.90 kb | 719 bp | 1.8% |

*DMRscaler* iteratively calls DMR-like regions using windows of increasing size while integrating the results of each iteration into the next layer of DMRs. While the top-most layer is the primary output of *DMRscaler*, this procedure produces a nested hierarchy of DMRs when considering the list of results across all layers that allows for a nuanced view of the differential methylation architecture. In Figure 3B, the top layers of this hierarchy within the X-chromosome are shown as the DMRs called at layer 3 are consolidated in DMRs in layer 4, and then those DMRs in layer 4 are consolidated into a single DMR in layer 5 spanning the entire X-chromosome (Figure 3B).

While the entirety of the X-chromosome can be considered a differentially methylated feature, it has been well established that there is a small subset of genes on the X-chromosome that escape X-inactivation [48]. We note that there are regions within the hierarchical structure of the DMR on the X-chromosome that only consolidate at a higher layer (Figure 3B) and span regions that are known to escape X-inactivation and therefore do not have significant differential methylation between the sexes. The expectation when comparing methylation between females and males is that X-inactivation would result in differential methylation between sexes, with hypermethylation and to a lesser extent hypomethylation across the entire X-chromosome in females compared to males [49]. Therefore, regions where the Δβ between the two groups is at or near zero represent regions that escape X-inactivation due to a relative lack of differential methylation at this site. An example of one such region is the gap of two DMRs that persists until the final integration between layer 4 and layer 5 which occurs at chrX: 71,482,842 - 71,498,442 which corresponds to the CpG rich promoter of *RPS4X* (Figure 3C, S11), a gene known to escape X-inactivation [50]. This example demonstrates that while the top layer DMR, which spans the X-chromosome, correlates most intuitively with the phenomenon of X-inactivation, exploring the hierarchical structure of complex DMRs reveals biologically meaningful structure.

The complex hierarchical relation of DMRs within the X-chromosome contrasts with the DMRs of the autosomal chromosomes. DMRs on the autosome show little to no branching, which implies that these DMRs are stable at each iteration of the algorithm (Additional File 1: Figure S8). A genome view of one such DMR at chr9: 84,302,344-84,304,983 highlights this stability, where a feature identified as a DMR at the first layer of the algorithm is stable through each subsequent iteration (Figure 3D, S11). The gene *TLE1* overlaps this DMR and has previously been identified as an autosomal gene that is differentially methylated between males and females [51, 52].

The results of the differential methylation analysis between sexes highlights the utility of *DMRscaler* in identifying differential methylation features that exist at dramatically different scales in real data. This ability distinguishes *DMRscaler* from existing methods which either are unable to identify larger DMRs while preserving the stability of smaller DMRs, as in *DMRcate* and *comb-p*, or tend to fragment larger DMR into many smaller features, as in *bumphunter*. A brief analysis of the hierarchical structure that results from *DMRscaler*’s layer merging mechanism also suggests that *DMRscaler* captures biologically meaningful structure within a DMR, such as escape from X-inactivation. This ability to represent DMR structure more completely suggests *DMRscaler* as a potentially valuable tool for exploring the interactions between features of epigenetic regulation at different scales.

### Rare chromatin modifier syndromes contain regions of differential methylation spanning gene clusters critical for development

Next we analyzed DNA methylation datasets from several rare diseases of chromatin modifier genes to see whether *DMRscaler* revealed novel DMR features that might otherwise be missed by existing methods. Except where stated otherwise, in the following sections DMRs are assumed to be those in the most inclusive top layer, Layer 5, which is built using all lower layers and is meant to be the most accurate representation of DMR features.

First, we compared the DNA methylation profile from fibroblasts from *KAT6A* patients to control samples. This analysis consisted of 20 samples, with 8 patients and 12 controls (Additional File 1: Table S2). All patients were previously reported by Kennedy, et al [7]. In our analysis, *DMRscaler* identified 575 unique DMRs with a median width of 2.19 kb (IQR: 278 bp -90.9 kb), resulting in a total genomic coverage of 3.5% (99.88 Mb) (Additional File 1: Table S11, Figure S12). Over the *HOXB* gene cluster two unique DMRs were identified. The first DMR spans *HOXB3, HOXB4, HOXB5,* and *HOXB6* and is hypomethylated in KAT6A patients relative to controls. The second spans *HOXB9* and is hypermethylated in KAT6A patients relative to controls. A more detailed look shows that the DMR spanning *HOXB9* is composed of two lower layer DMRs that were consolidated in the 16 Adjacent CpG layer, with the *HOXB9* promoter being flanked but not captured in the lower layer DMRs (Figure 4A,B, Additional File 1: Figure S13). The relatively large coverage of the genome by DMRs is driven primarily by multi-megabase scale DMRs identified spanning relatively gene sparse regions (Figure 4C,D, Additional File 1: Figure S13).

**Figure 4:**
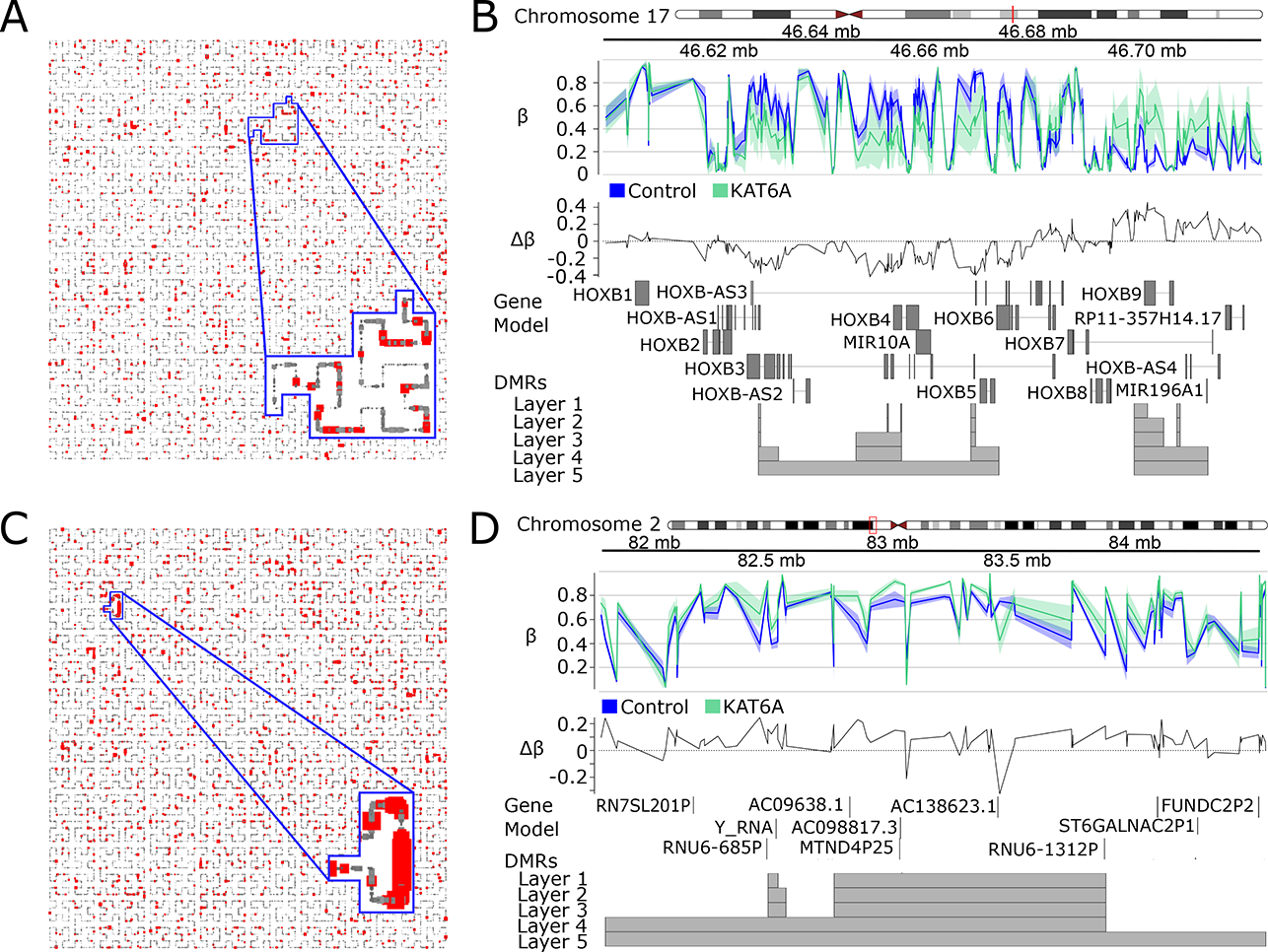
Differential Methylation Analysis in KAT6A Syndrome. (A) Hilbert curve of CpGs from chr17, outlined is the region corresponding to Chr17:46.59-46.73Mb, the *HOXB* cluster. CpGs with FDR < 0.1 are highlighted red. Point size is scaled to maximum significance value. (B) Chr17:46.59-46.73Mb. *HOXB* cluster. GVIZ track stack plot. Top track shows mean β value per group, next track shows Δβ, where Δβ = β_KAT6A_ - β_Control_. Below the gene model track is the DMR track, highlighting the regions called as a DMR at each result layer from *DMRscaler*. (C) Chr2:81.5-84.5 Mb. Design same as 4A. **(D)** Chr2:81.5-84.5 Mb. Tracks same as 4B.

Weaver syndrome (MIM# 277590), is an rare overgrowth disorder that is caused by de novo mutations in *EZH2*, a histone methyltransferase. Comparing Weaver syndrome patient samples to controls, *DMRscaler* identified 485 unique DMRs with a median width of 1.06 kb (IQR: 378 bp - 10.2 kb). These regions comprised a total of 0.34% (9.66 Mb) of the genome (Additional File 1: Table S12, Figure S14).

Over the *HOXA* gene cluster, *DMRscaler* identified two distinct DMRs associated with Weaver Syndrome. The first spans *HOXA1-HOXA6.* The structure of this DMR is complex, with *HOXA1, HOXA2*, and *HOXA4* overlapping CpGs modestly hypermethylated in Weaver syndrome, and *HOX5* and the last two exons of *HOX6* overlapping CpGs with a greater magnitude of hypomethylation. The second DMR spans *HOXA9-HOXA11, HOXA13, HOTTIP,* and the nearby gene *EVX1*. This second DMR is generally weakly hypermethylated in Weaver Syndrome, with a small but significant region of hypomethylation just upstream of *HOXA11* (Figure 5 A,B, S15).

**Figure 5:**
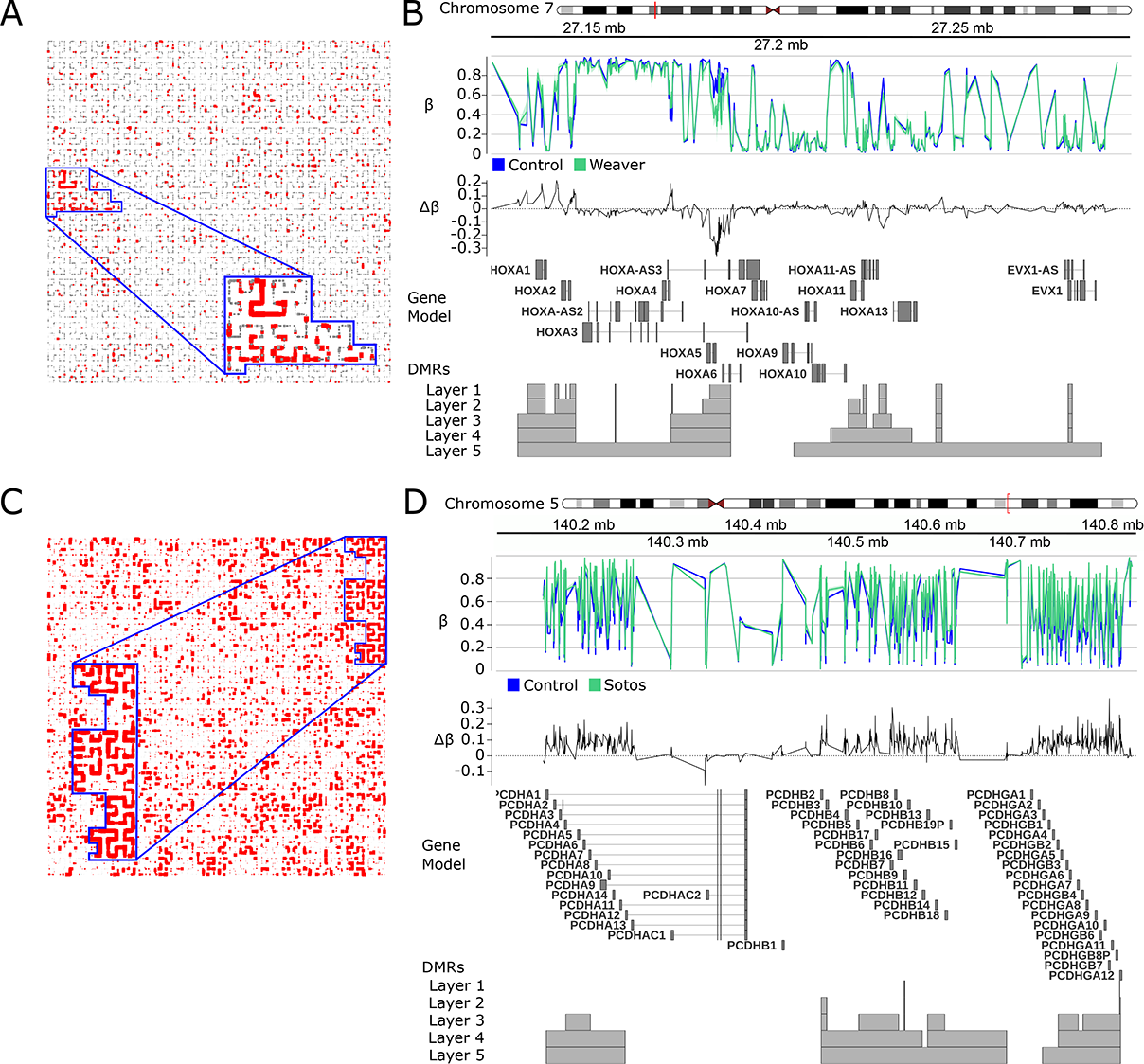
Differential Methylation Analysis in Weaver (A,B,C) and Sotos Syndrome (D,E,F). (A) Hilbert curve of CpGs from chr7, outlined is the region corresponding to Chr7:27.1-27.3Mb, the *HOXA* cluster. CpGs with FDR < 0.1 are highlighted red. Point size is scaled to maximum significance value. (B) Chr7:27.1-27.3Mb. *HOXA* cluster. GVIZ track stack plot. Top track shows mean β value per group, next track shows Δβ, which is β value relative to mean female β value. Below the gene model track is the DMR track, highlighting the regions called as a DMR at each result layer from *DMRscaler*. (C) Chr5:140.1-140.8Mb over the *PCDH* clusters. Design same as 4A. (D) .Chr5:140.1-140.8Mb over the *PCDH* clusters. Tracks same as 4B.

Finally, we also analyzed Sotos Syndrome (MIM# 117550), an overgrowth syndrome caused by truncating and missense mutation in the nuclear receptor binding SET domain protein 1 (*NSD1)* gene [53]. Analysis with DMRscaler identified 757 unique DMRs with a median width of 1.87 kb (IQR: 617 bp - 25.3 kb), covering 23.1 Mb of the genome (0.8%) (Additional File 1: Table S13, Figure S16). We identified four unique DMRs that span gene clusters related to protocadherins, large transmembrane proteins that are critical for a diverse range of processes ranging from cell-signaling to dendritic arborization [54]. These DMRs caused by mutations in *NSD1* cover the neighboring *Protocadherin* (*PCDH*) gene cluster *PCDHA*, *PCDHB*, and *PCDHGB* (Figure 5D,E, S15). One DMR spans the first exons of *PCDHA1-PCDHA12*, another spans from *PCDHB2* to *PCDHB19P*, and a third covers the first exons of *PCDHGA3-PCDHGA12* and *PCDHB1-PCDHB8P*, and finally a smaller DMR covers the first exon of *PCDHGC3*. All of the DMRs covering these *PCDH* clusters are hypermethylated in Sotos Syndrome relative to controls, though it is notable that the β values of CpGs across these clusters are highly variable reflecting an example of the neighboring CpG heterogeneity described earlier (Figure 5D, S15, S4, S5).

These results in rare chromatin modifier syndromes highlight *DMRscaler’s* utility in identifying patterns of differential methylation that exist over broader genomic features such as gene clusters.

These results in rare chromatin modifier syndromes highlight *DMRscaler’s* utility in identifying patterns of differential methylation that exist over broader genomic features such as gene clusters.

### Analysis of overlapping regions of differential methylation

Following analysis of each syndrome individually, we asked whether there was evidence of shared regions differentially methylated between KAT6A, Sotos, and Weaver syndrome. Between KAT6A and Sotos syndrome, we identified 21 regions with overlapping DMRs (8.2% of total DMRs for KAT6A, 3.0% of total DMRs for Sotos), and 28 genes overlapped by some DMR in both syndromes (11.4% of genes overlapping a DMR in KAT6A syndrome, 4.1% of genes overlapping a DMR in Sotos syndrome). Between KAT6A and Weaver syndrome, we identified 22 regions with overlapping DMRs (8.6% of total DMRs for KAT6A syndrome, 4.7% of total DMRs for Weaver syndrome), and 37 genes overlapped by some DMR in both syndromes (15.0% of genes overlapping a DMR in KAT6A syndrome, 8.3% of genes overlapping a DMR in Weaver syndrome). Between the two growth disorders, Sotos and Weaver syndrome, we identified 55 regions (8.1% of total DMRs for Sotos, 13.6% of total DMRs for Weaver) and 65 genes overlapped by some DMR in both syndromes (9.4% of genes overlapping a DMR in Sotos, 14.6% of genes overlapping a DMR in Weaver) (Additional File 6: Table S8).

To test the significance of the overlap we tested the odds ratio (OR) of overlap between each pair of syndromes. To simplify the analysis and make the measure of the odds ratio closer in form to the *DMRscaler* method, we only used counts of measured CpGs (See methods for details). Essentially, the odds ratio tests whether there is enrichment of CpGs that are in DMRs in one syndrome in the set of CpGs found in DMRs in the other syndrome being compared. OR with a confidence interval (CI) overlapping 1 suggests no enrichment, closer to 0 or further from 1 indicates greater enrichment. The raw overlap counts used to calculate the OR are in (Additional File 1: Table S9) and the odds ratios are reported in (Additional File 1: Table S10). The highest odds ratio was between Sotos and Weaver with OR = 9.39 (95% CI: 8.11-10.86) in Layer 1. The OR between Sotos and Weaver drops to OR = 1.75 (95% CI: 1.59-1.92) in Layer 5, which might be due to decreased similarity over broader DMR features than those in Layer 1 which have the highest density of significantly differentially methylated CpGs. In contrast, the OR between KAT6A and Weaver is relatively consistent between Layer 1 and Layer 5, with OR = 3.71 (95% CI: 2.42-5.66) and OR = 4.32 (95% CI: 3.78-4.93). The OR between KAT6A and Sotos follows a similar decrease to that between Sotos and Weaver, with OR = 3.9 (95% CI: 2.95-5.15) in Layer1 and OR = 1.48 (95% CI: 1.24-1.76) in Layer 5 (Additional File 1: Table S10). This implies DMRs of each of the three syndromes analyzed here are enriched in CpGs in DMRs in both of the other syndromes tested here, even though the number of features with overlap is modest.

One region of overlap between Sotos and Weaver syndrome was a region overlapping *INS,* and *INS-IGF2* proximal to *IGF2*. This region stood out as a region implicated in another growth disorder, Beckwith-Wiedemann syndrome (BWS) [55]. The DMR called for Sotos syndrome is just upstream of *IGF2* and overlaps *INS* and *INS-IGF2*, with sites of moderate effect hypomethylation (Δβ > 0.2) (Additional File 1: Figure S17A). The DMR called for Weaver syndrome overlaps the IGF2 gene, though it spans further 3’ than the Sotos syndrome DMR. This DMR in Weaver syndrome was composed of sites with a small effect size (Δβ ∼ 0.05 ), with hypermethylation over a region of the *IGF2* gene body and hypomethylation further upstream overlapping the *INS* and *INS-IGF2* genes. Upstream of *IGF2* overlapping *INS* and *INS-IGF2* the pattern of hypomethylation in Sotos and Weaver syndrome was consistent (Additional File 1: Figure S17B, Additional File 6: Table S8).

We did not identify any regions of direct overlap in all three syndromes, however there were 13 genes overlapping some DMR in all three syndromes. These were *TFAP2E, SPON2, PCDHGA1, PCDHGA2, PCDHGA3, PCDHGA8, PCDHGA10, PCDHGB7, PCDHGA12, PCDHGC3, GATA4, HOXC6, HOXC4*. The *PCDHG* cluster genes, previously discussed in context of Sotos syndrome alone, are worth noting as they are involved in neural development. The *PCDHG* cluster is broadly hypermethylated in Sotos, as noted earlier (Figure 5C,D), with a second narrower region of hypermethylation at the 5’ end of *PCDHGC3*. In KAT6A syndrome, there is a DMR intergenic to most of the *PCDHG* genes and at the 5’ end of *PCDGC3*, and in Weaver syndrome there is a DMR with minor hypomethylation at the shared 3’ end of *PCDHG* genes (Additional File 1: Figure S18).

The overlapping DMRs and genes with overlapping DMRs between KAT6A, Sotos, and Weaver syndrome reveals a number of shared regions of differential methylation and shared genes with patterns of differential methylation. CpGs in DMRs in any one syndrome are enriched in CpGs in DMRs of either of the other syndromes across all three pairs of syndromes, KAT6A-Sotos, KAT6A-Weaver, Sotos-Weaver, as measured by the odds ratio. However, these data are derived from different cell types (fibroblast vs blood) and exhibit cell-type specific changes in addition to those caused by the genetic mutation. Together these results suggest that while each syndrome has a distinct profile of differential methylation, there is also significant overlap in regions mirroring shared phenotypic features.

## DISCUSSION

The key development of our new method, *DMRScaler*, is a substantial improvement in the ability to accurately identify the size of DMRs across the full range of epigenetic scale. Whole genome bisulfite sequencing is technically and analytically challenging and remains prohibitively expensive for routine use. The majority of real-world methylation data is in the form of reduced representation platforms that query CpGs in sites that are likely to play a role in gene regulation, such as known enhancers and transcriptional start sites. While the distance between the sites are variable on an array, our sex chromsome results demonstrate the ability of our method to call established DMRs that vary in size.

Differential methylation analysis between sexes performed with *DMRscaler* showed our algorithm could handle the full range of DMR features present. Looking at the DMRs between XX and XY individuals, *DMRscaler* was able to identify a small DMR 2.6 kb in length overlapping the autosomal gene *TLE1* that had previously been identified as differentially methylated between the sexes [51], while also consolidating the differential methylation of the X-chromosome into a single DMR 152.11 Mb in length, spanning 98% of the total length of the chromosome.

Additionally, *DMRscaler* provides the means for a hierarchical definition of a DMR that is built through the iterative procedure of merging layers built from increasing window sizes. A deeper analysis of the DMR spanning the X-chromosome showed that a late consolidating region, where a gap remained until the integration of the 64 CpG layer, overlapped the *RPS4X* gene which is known to escape X-inactivation [50], which is concordant with data showing that regions escaping X-inactivation should have a similar epigenetic landscape between sexes. These results show how *DMRscaler* provides an intuitive representation of DMRs and provides a mechanism for a hierarchical definition of DMRs that can be used to investigate the structure of the methylation landscape across larger epigenomic features. Together these behaviors of the method allow greater flexibility and more meaningful interpretation of results in analyses than existing methods.

Finally, given our primary interest in leveraging this method for the smaller sample sizes in rare-disease studies, we tested *DMRscaler* on datasets from patients with rare chromatin modifier syndromes. Specimens that harbor known pathogenic mutations in chromatin modifier genes often display regional changes to epigenetic features, such as DNA methylation state [9, 10]. Our study also explored three syndromes that are caused by pathogenic mutations in genes that directly control histone modifications.

Arboleda-Tham Syndrome (MIM# 616268) syndrome is a genetic syndrome caused by mutations in the Lysine (K) acetyltransferase *KAT6A* characterized by global developmental delay, intellectual disability, speech delay or absence and phenotypes of variable expressivity such as congenital heart defects and gastrointestinal anomalies [7, 56]. KAT6A acetylates histones K3K9, H3K14, and H3K23 [57–59], but the genomic regions affected by KAT6A have not been comprehensively studied. Previously, deletion of *KAT6A* in model organisms has identified the *HOX* genes, including the *HOXB* cluster, as regulatory targets of KAT6A [58, 60]. Two DMRs identified here were identified by *DMRscaler* spanning multiple genes of the *HOXB* cluster (Figure 3), which encompass 4 genes (*HOXB3 - HOXB6)* found in a *KAT6A* knockout mouse model to have shifted domains of expression resulting in homeotic transformation of the axial skeleton [58]. The ability to highlight the extent of differential methylation beyond a single gene provides further context into the epigenetic change that occurs in *KAT6A* syndrome.

One limitation of our study is that cases are generally younger in age than the control groups. Previous studies have identified global hypomethylation as associated with aging [61, 62]. For regions that are largely hypermethylated in *KAT6A* Syndrome patients relative to controls, we cannot exclude this as a potential confounding factor in our analysis.

Weaver syndrome and Sotos syndrome are rare overgrowth syndromes that can be difficult to distinguish without sequencing. They are caused by mutations in *EZH2* [63, 64] and *NSD1* gene [53], respectively. Despite their common clinical phenotype of overgrowth, the regions of the genome that are identified as differentially methylated largely diverge between these two syndromes suggest distinct pathways to a common and complex phenotype. For Weaver Syndrome, *DMRscaler* identified differential methylation over the *HOXA* cluster genes in Weaver syndrome relative to controls. Genome-wide mapping of EZH2 binding domains shows EZH2 binds the *HOXA* cluster [65] and EZH2 overexpression in mantle cell lymphoma has been associated with hypermethylation over the *HOXA* cluster [66]. The key improvement is that rather than highlighting individual genes [9] as differentially methylated, *DMRscaler* is able to demonstrate the modular nature of the genetic regulation by highlighting the non-random spatial relation of these features as a pair of DMRs spanning several genes each.

Additionally, *DMRscaler* identified a novel finding in the neighboring *PCDHA, PCDHB, and PCDHG* clusters as broadly hypermethylated between Sotos syndrome patients relative to controls. The protocadherin family genes are critical in cell-cell adhesion and involved in the complex patterning of neural circuitry [54]. These same genes in the *PCDHGA/B* cluster were also identified as hypermethylated in Down syndrome human cortex relative to control cortex tissue [67]. From these results we can hypothesize that misregulation of the *PCDH* clusters in brain development contributes to the neurodevelopmental phenotype of Sotos syndrome, however this remains to be tested.

Notably, we observed that between the two overgrowth syndromes, Sotos and Weaver syndrome, the *IGFR2* region including the *INS* and *INS-IGFR2* genes was similarly differentially methylated. Loss of normal imprinting regulation of *IGFR2* has been implicated in another overgrowth syndrome, Beckwith-Wiedemann Syndrome (BWS) [55]. Whether this common difference in DNA methylation proximal to the *IGFR2* locus represents an epigenetic contributor to the overgrowth phenotype or is a consequence of the overgrowth phenotype is worth further investigation.

While empirically *DMRscaler* works well to identify regions of differential methylation across a large dynamic range, improvements in the statistical framework for integrating regions across layers of DMRs called with increasing window sizes could allow for extension of these approaches in other types of functional epigenomic datasets.

## CONCLUSION

Here we have shown that *DMRscaler* is flexible yet robust in describing the scale of DMR features from the local scale of individual promoters and CpG sites, to the DMR features that represent chromosome level differences in methylation. All of the analyses described were run using a shared parameter set for *DMRscaler*, which highlights the utility to researchers who seek to explore these higher order epigenetic features while also describing the local changes with known biological implication, such as changes in methylation overlapping the promoter of a gene. Importantly, *DMRscaler* serves as a proof of principle. The idea that important epigenetic features exist beyond the scale of a single gene is not new to most researchers. However, existing methods for DNA methylation analysis do not capture this knowledge. Here *DMRscaler* proves that it is possible to computationally capture this intuition, and in doing so reveal novel biological insights. *DMRscaler* serves as a framework for analysis of epigenetic information in a manner that accurately identifies feature size across the full range of epigenomic scale.

### ABBREVIATIONS

TF: Transcription factor
PRD: Polycomb repressive domain
TAD: Topologically associated domain
DNAme: DNA methylation
CpG: Cytosine guinine dinucleotide
VUS: Variant of unknown significance
DMR: Differentially methylated region
FDR: False discovery rate
QC: Quality control
TP: True positive
FN: False negative
FP: False positive
P: Precision
R: Recall
AUCPR: Area under precision recall curve

## SUPPLEMENTAL INFORMATION DESCRIPTION

Additional File 1 (pdf):

Supplemental Figures S1-18; Supplementary Tables S1-S3, S9-S13

Additional File 2: Supplemental Table 4: KAT6A DMRs (xlsx)

Additional File 3: Supplemental Table 5 (xlsx) : Differential Sex Analysis DMRs

Additional File 4: Supplemental Table 6 (xlsx) : Weaver Syndrome DMRs

Additional File 5: Supplemental Table 7 (xlsx) : Sotos Syndrome DMRs

Additional File 6: Supplemental Table 8 (xlsx) : Syndrome Overlaps

## DECLARATIONS

### Ethics approval and consent to participate

The data used in this study was approved by the UCLA Institutional Review Board, study number IRB#11-001087.

### Consent for publication

Not applicable

### Availability of data and materials

Project Name: DMRscaler. Project home page: https://github.com/leroybondhus/DMRscaler. Operating system(s): Platform independent. Programming language: R. Other requirements: R version 4.1.0 or higher [68]. License: MIT. Any restrictions to use by non-academics: None. The code for the simulation analysis is available at https://github.com/leroybondhus/dmrscaler_simulation. The code for the real data analysis is available at https://github.com/leroybondhus/dmrscaler_real_data. Datasets from Sotos and Weaver Syndrome can be found with GEO accession: GSE74432 at https://www.ncbi.nlm.nih.gov/geo/query/acc.cgi?acc=GSE74432 Methods used for comparison can be found at DMRcate [45]: https://www.bioconductor.org/packages/release/bioc/html/DMRcate.html bumphunter [43] : https://www.bioconductor.org/packages/release/bioc/html/bumphunter.html comb-p [44] : https://github.com/brentp/combined-pvalues Packages used for visualization of data Hilbert Curves [69]: https://www.bioconductor.org/packages/release/bioc/html/HilbertCurve.html Genomic Range Plots [70] : http://bioconductor.org/packages/release/bioc/html/Gviz.html Network Plots [71] : https://cran.r-project.org/web/packages/networkD3/index.html

### Author’s Contributions

LB and VAA conceived of and designed the study. LB wrote code, analyzed data and made figures. AW wrote code for the simulation approach and analyzed data for the overlap analysis. VAA and LB interpreted the results and wrote the manuscript.

## Acknowledgements

We thank the members of the Arboleda lab for their helpful comments and feedback on the manuscript. We thank the UCLA Neuroscience Genomics Core for their support and expertise in processing our samples.

