## Additional File 1 for "*DMRscaler*: A Scale-Aware Method to Identify Regions of Differential DNA Methylation Spanning Basepair to Multi-Megabase Features"

*Corresponding Author

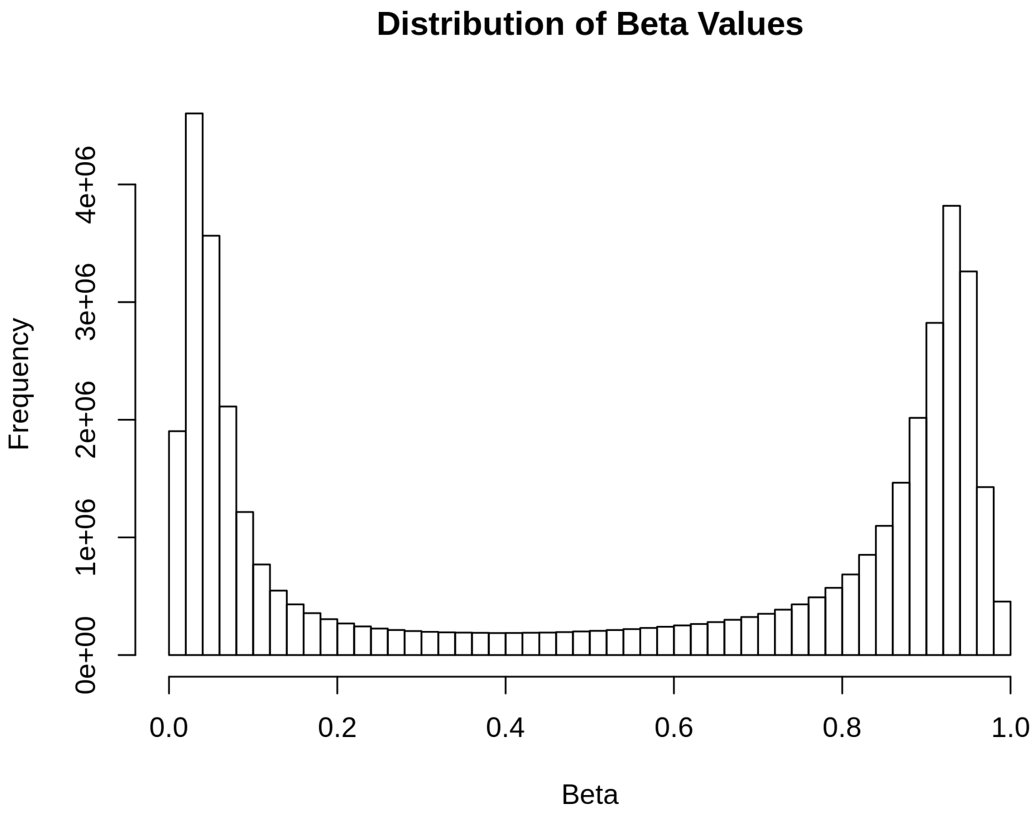


**Figure S1.** Beta Distribution of methylation data from Illumina Infinium Human Methylation450 Bead Chip 450 array. Beta value is the proportion of methylation at each CG site.

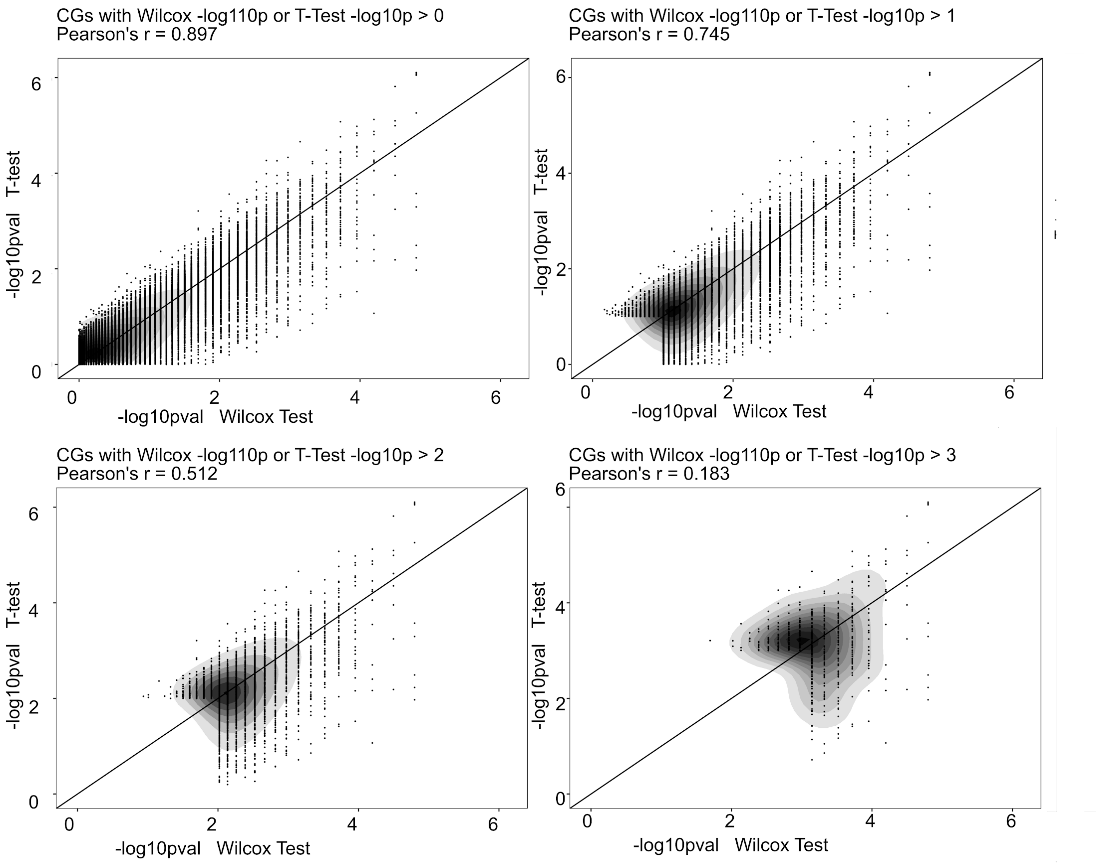


**Figure S2**: Wilcox - T-Test pearson's correlation (pearson's r) for each CG. From KAT6A data, case-control labels determine group.

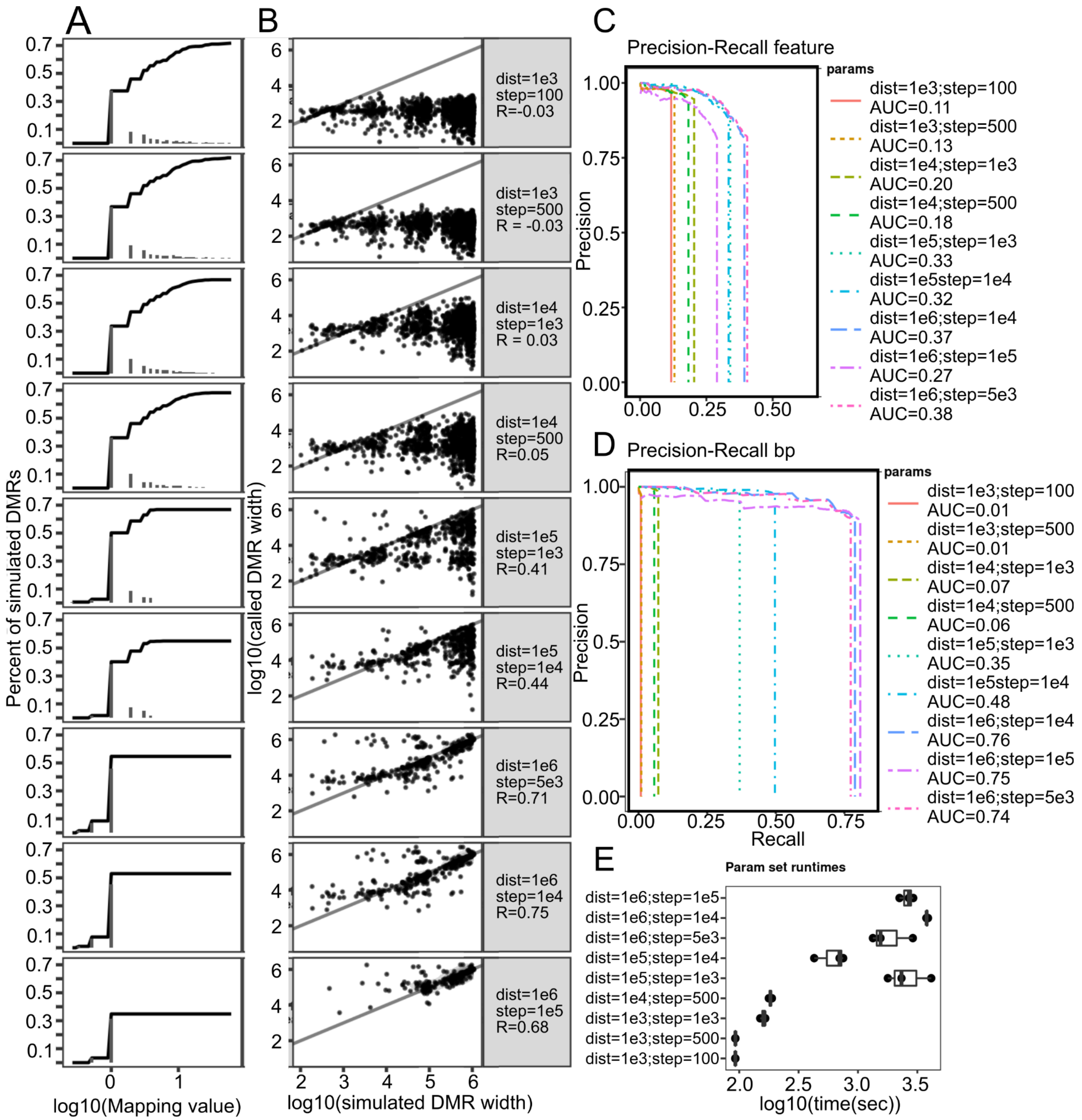


**Supplement Figure S3: comb-p parameter testing**: **(A)** Mapping Values plots. Log values > 0 imply multiple DMRs called per simulated DMR. Value < 0 imply multiple simulated DMRs overlap single called DMR. Value = 0 implies one DMR called per DMR simulated. The plotted line indicates the cumulative proportion of simulated DMRs up to the given mapping value. **(B)** Simulated DMR Widths v Called DMR Widths plotted on log10 scale.**(C)** Feature level precision-recall curves, see methods for details on calculation. **(D)** basepair level precision recall curves **(E)** time for each parameter set run on the simulated dataset across 3 runs.

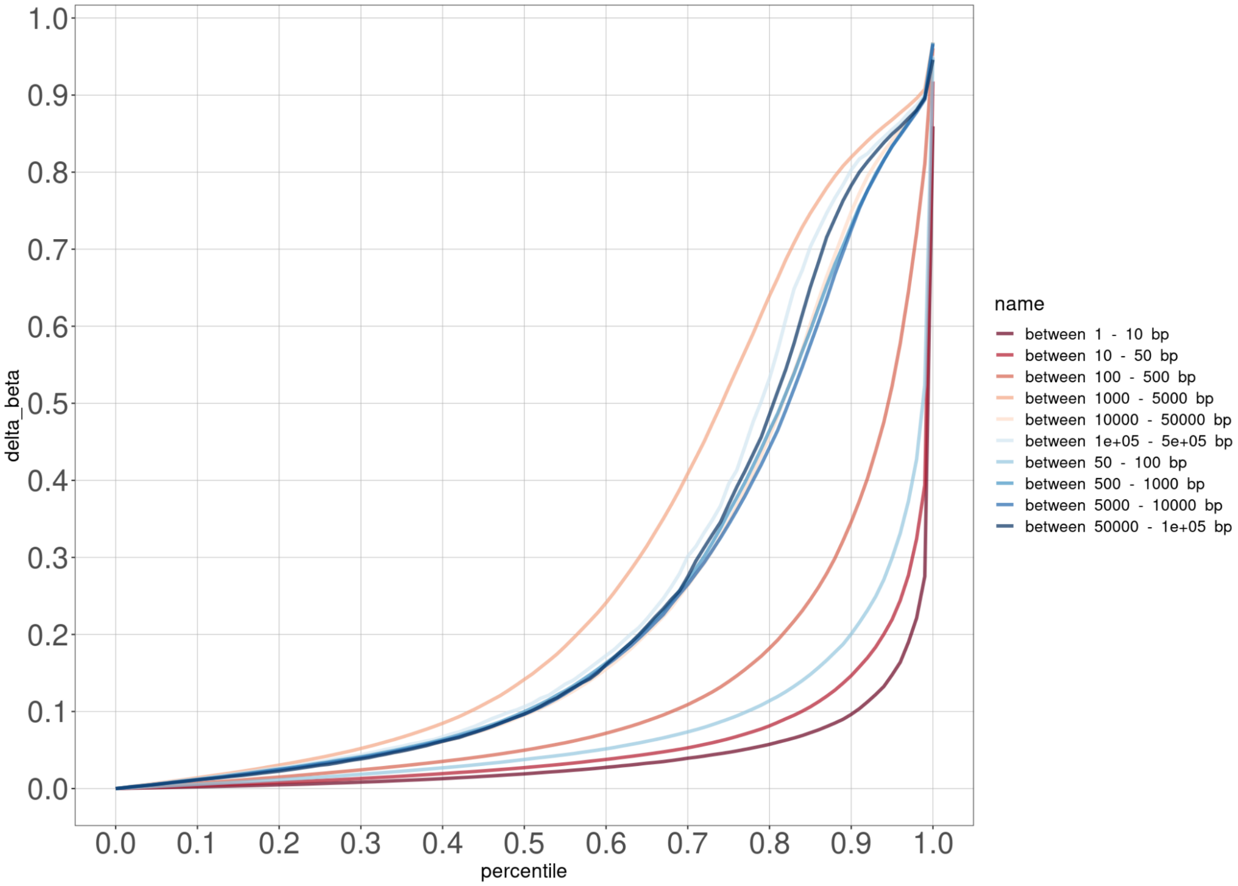


**Figure S4**: Cumulative Probability of difference in beta values between neighboring CG methylation. Y axis is the difference in Beta between neighboring CGs (CG_i and CG_i+1). E.g. 70% of adjacent measured CGs that are between 1000-5000 bp apart have a difference in beta value less than 0.40, said another way, 30% have a difference in beta greater than 0.4.

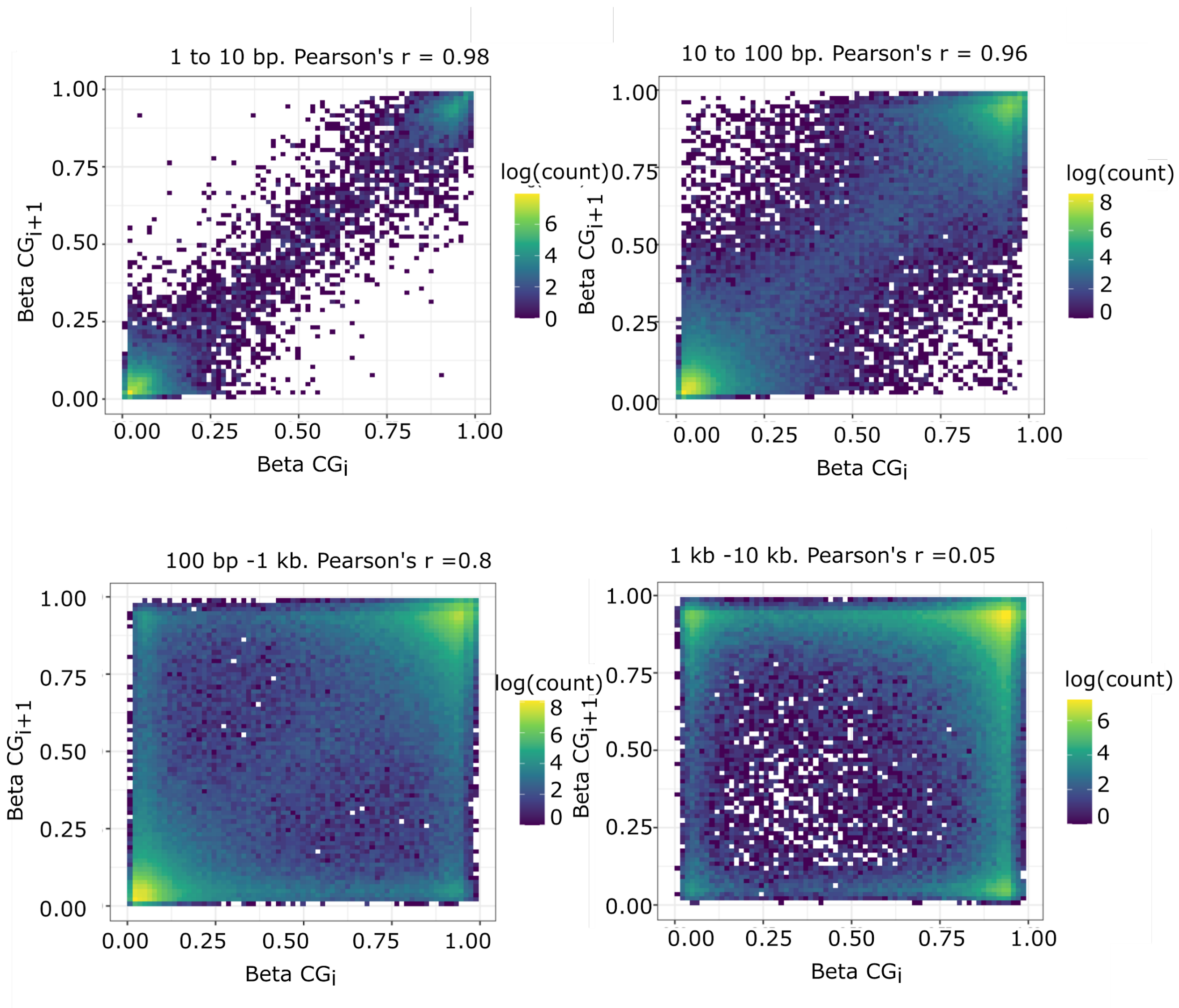


**Figure S5:** Neighboring CG methylation correlation. Plotted is beta value from 0-1, for CG_i on the x-axis and CG_i+1 on the y-axis. Neighboring CGs 1-10 bp apart and 10-100 bp apart have strong, but not perfect correlation (Pearson's r = 0.98, 0.96), those 100-1000 bp apart have modest correlation (Pearson's r = 0.8). At 1-10kb distance the correlation becomes weak (Pearson's r = 0.05).

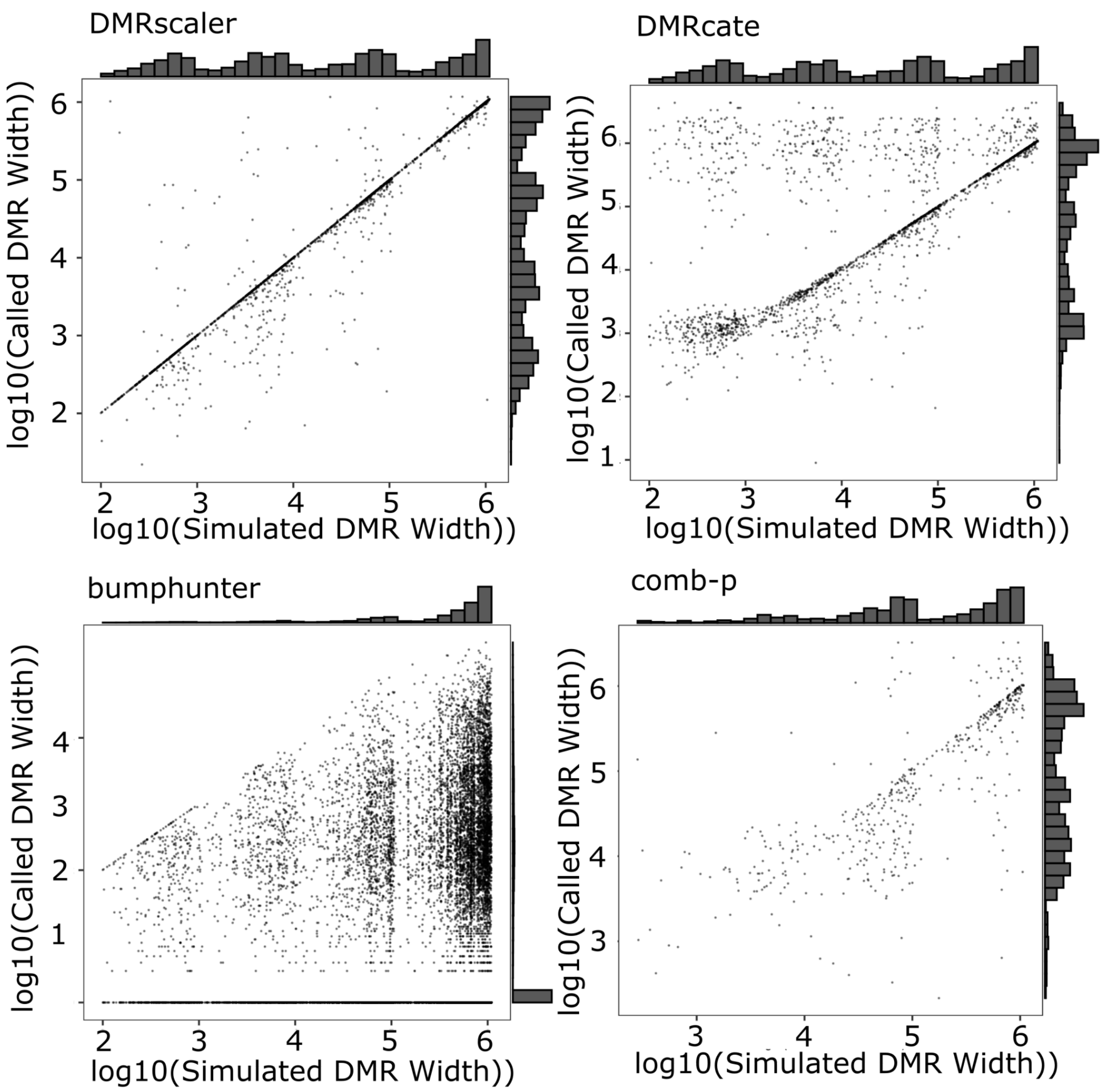


**Figure S6**. Simulated vs Called Widths with marginal density plots.

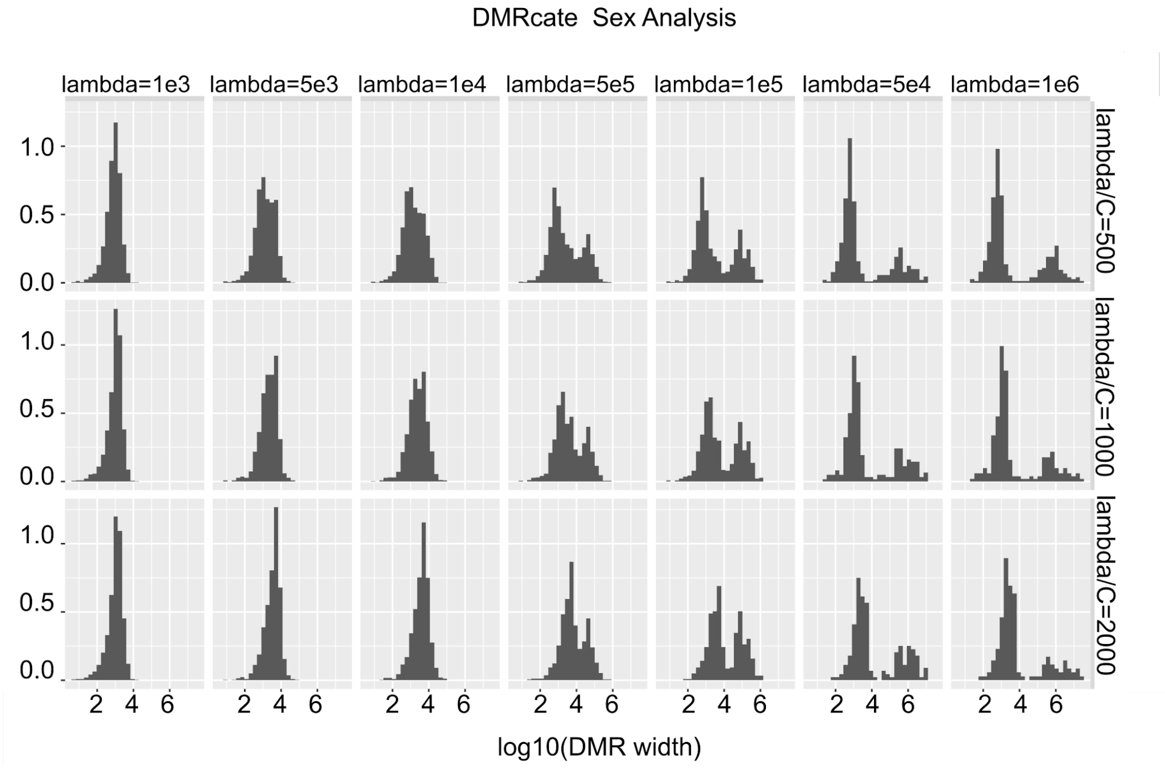


**Figure S7:** Testing variable lambda and C parameters on output from sex analysis for DMRcate. Only showing autosomal DMRs.

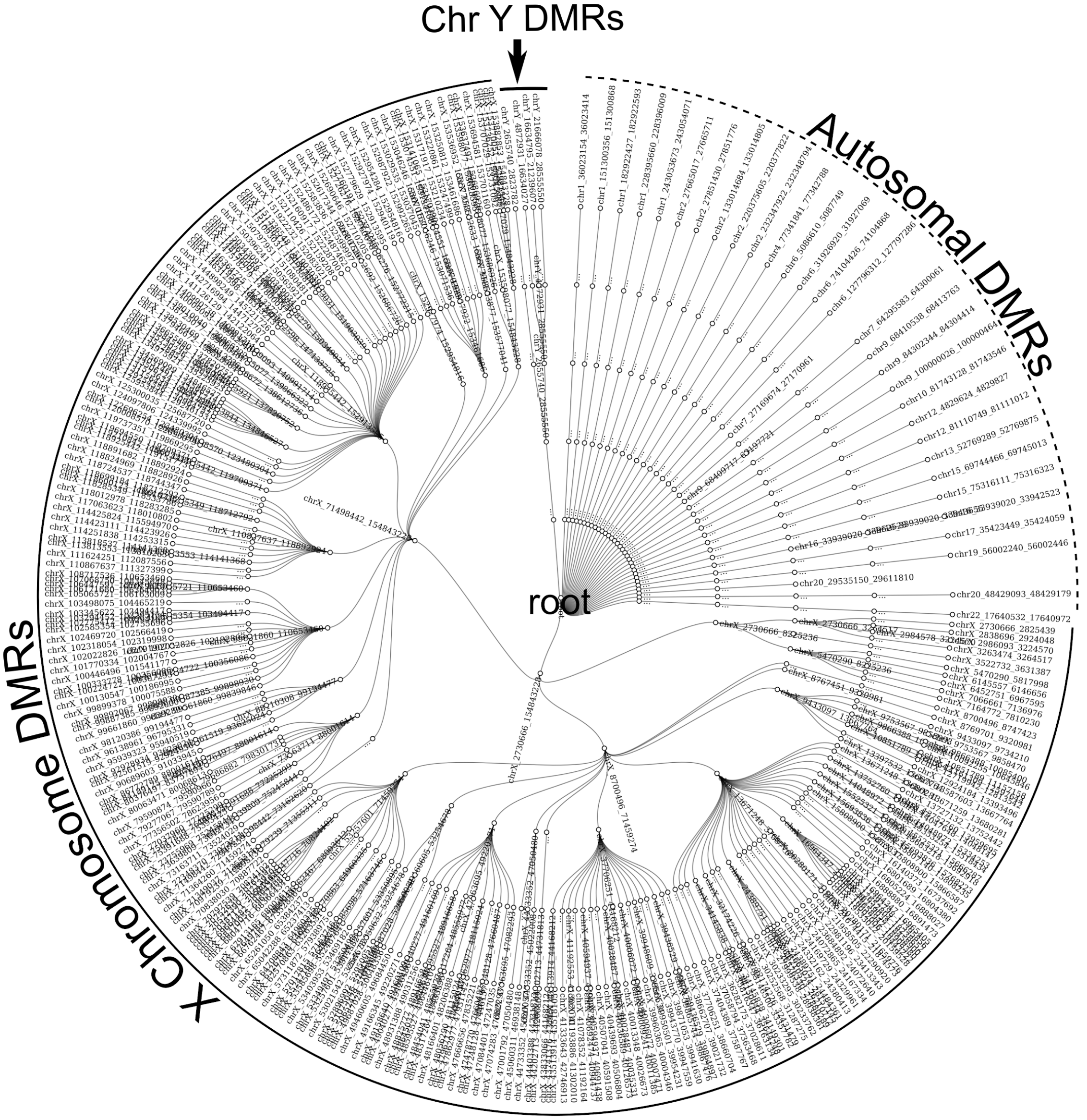


**Figure S8**. Radial Network Showing hierarchical structure of DMRs called across all layers of the DMRscaler algorithm from layer 1 (4 adjacent CG windows) at the edges nodes to layer 5 (64 CG windows) in the inner ring. Note, all are connected to the virtual root node which is only used for plotting purposes here. Each node is an individual DMR called. Coordinates for each DMR are printed at the earliest layer where that DMR appears. Unlabelled nodes are those that did not change from the previous layer.

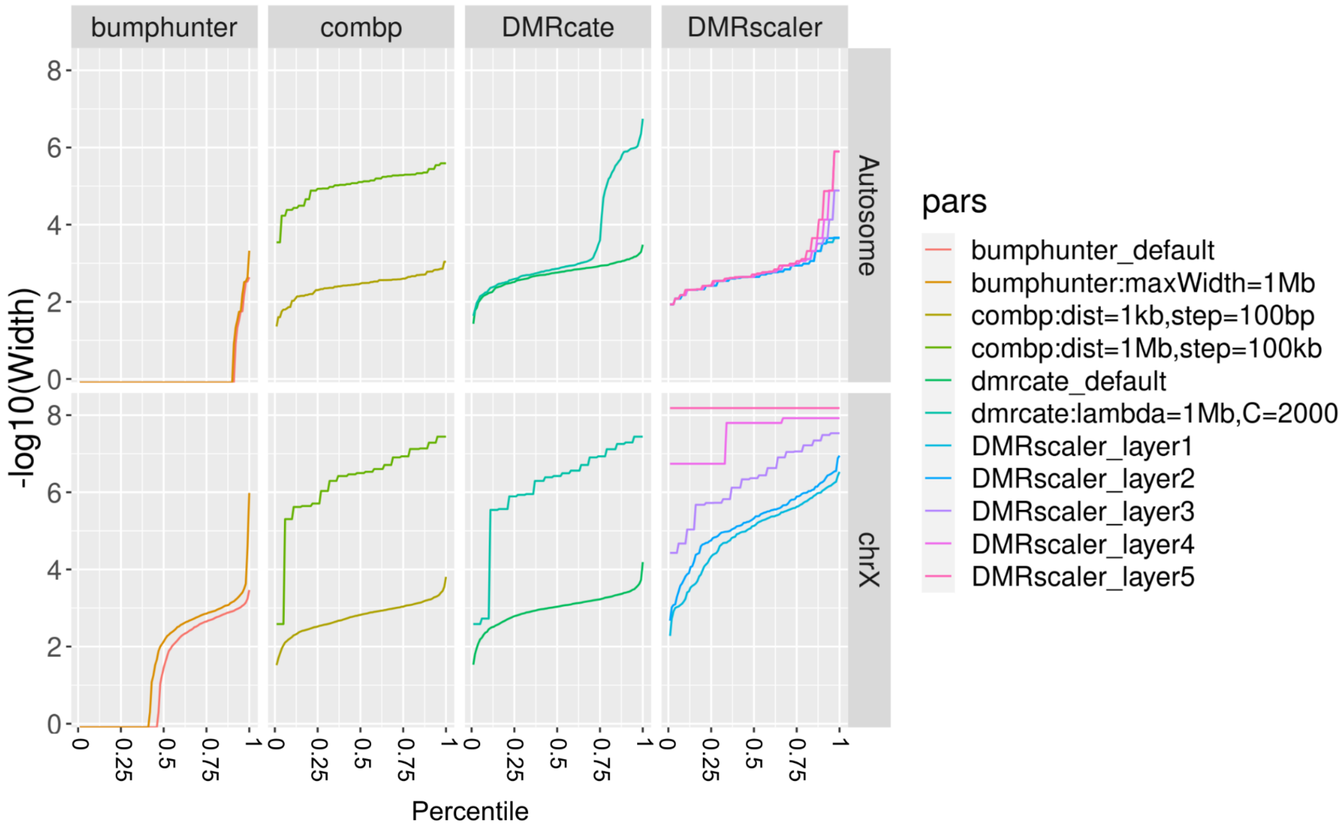


**Figure S9:** Percentile plot. DMRs Called by each method for sex analysis ordered by dmr width.

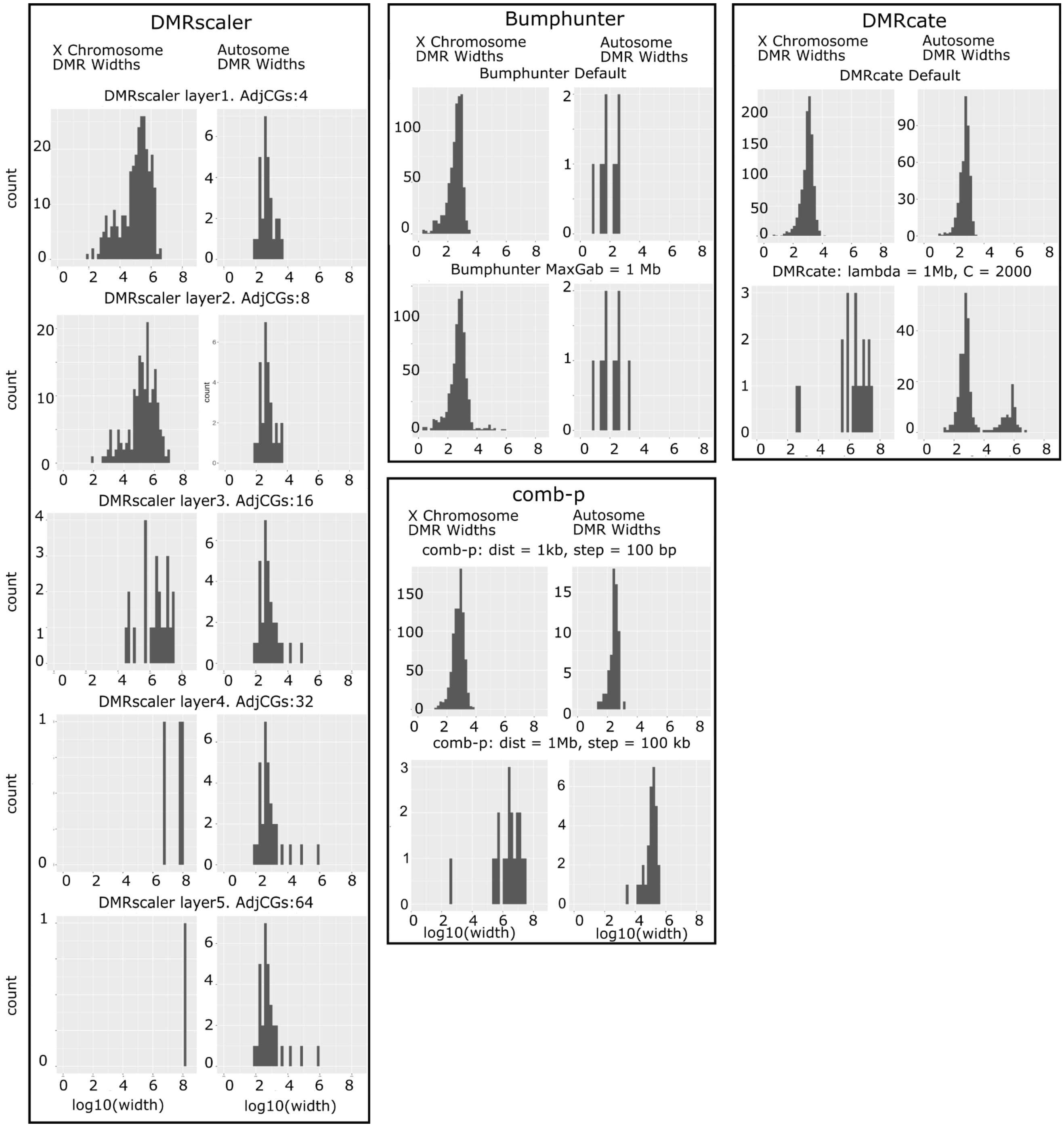


**Figure S10** : Distribution of DMR widths for each method called in XX vs XY sex chromosome analysis.

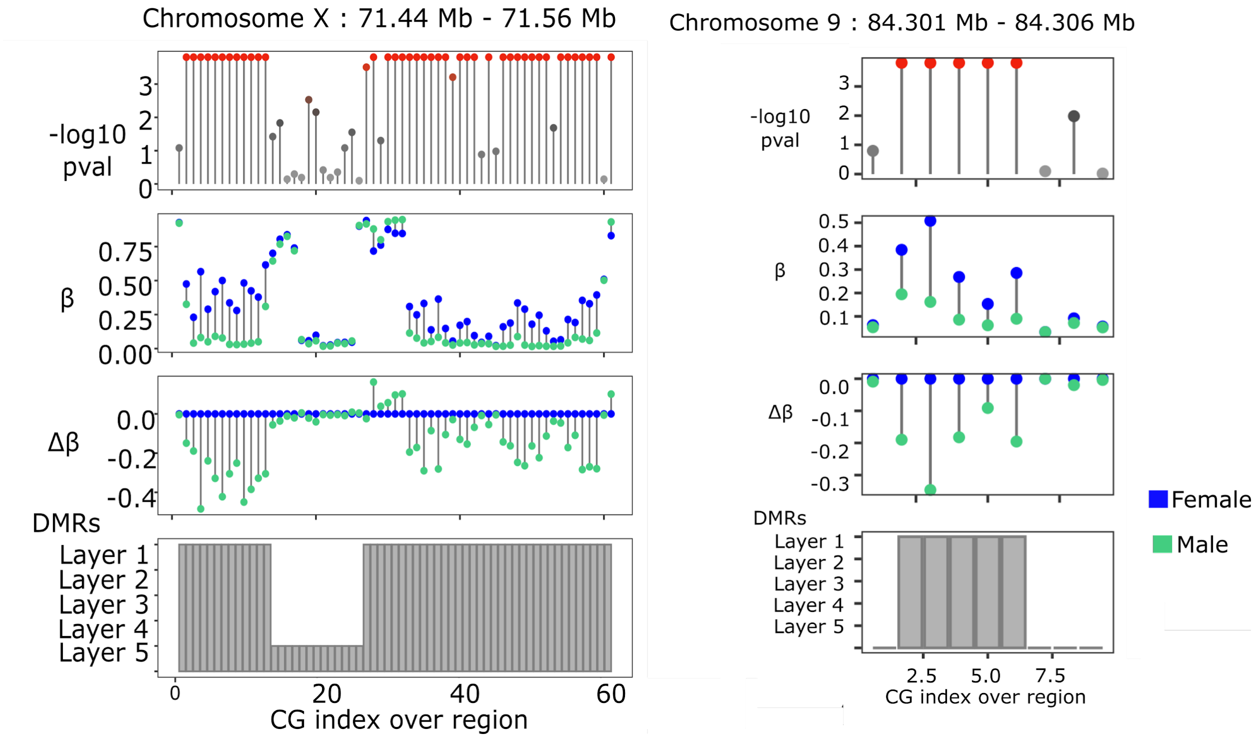


**Figure S11:** Sex analysis. Supplement to Figure 3C (left), 3D (right). Adjacency plot of CGs overlapping specified regions. Top panel is significance at individual CG level. Beta plot shows mean beta value for each group. Delta beta below shows mean beta value for each group relative to female values. Bottom plot shows in grey bars which layer a DMR was called in.

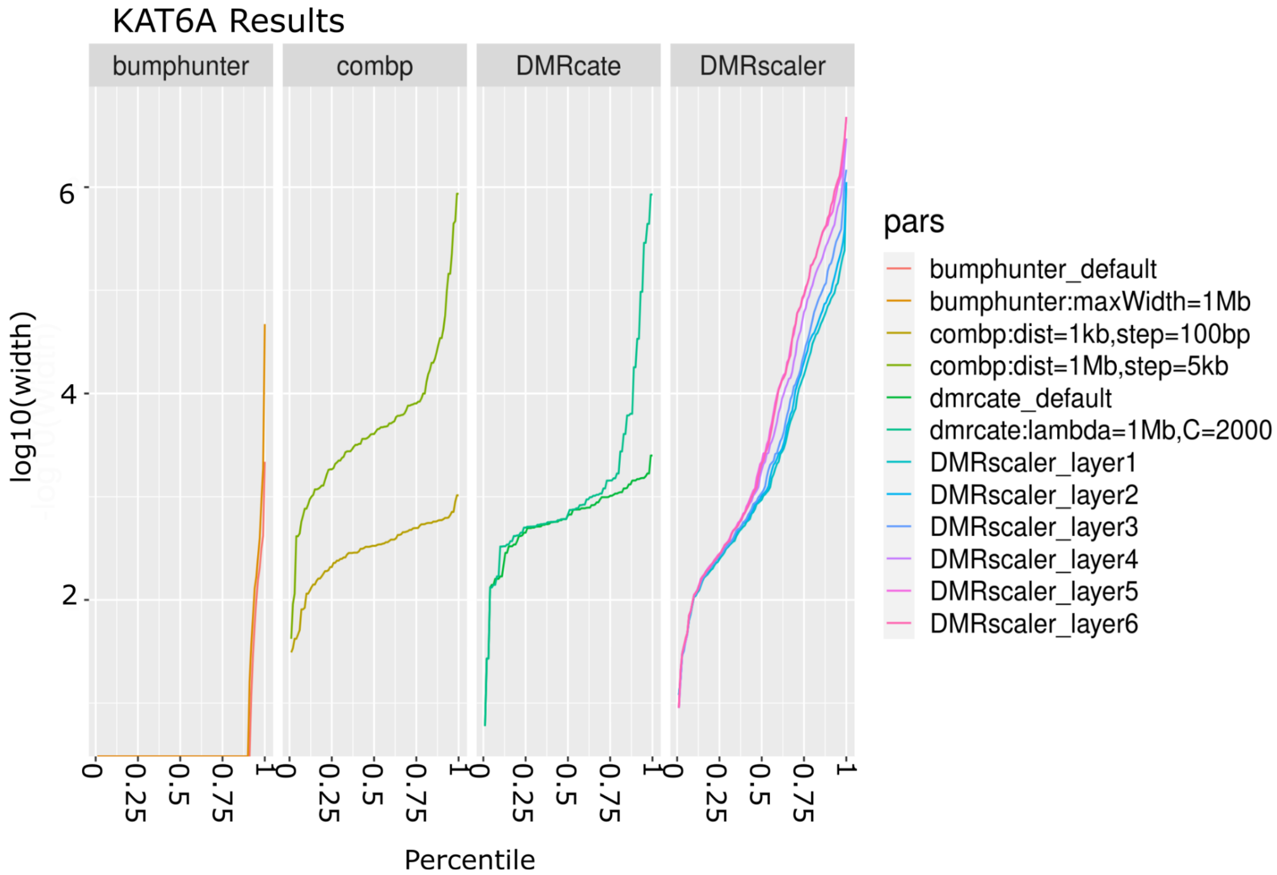


**Figure S12:** KAT6A Analysis: Called DMR size percentiles.

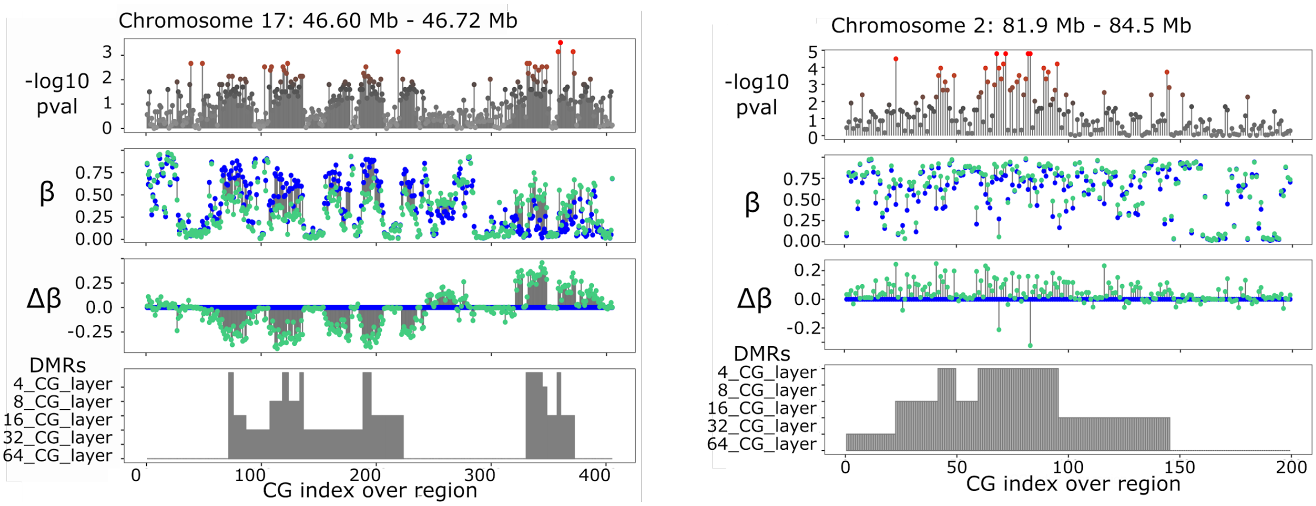


**Figure S13:** KAT6A analysis. Supplement to Figure 4B (left), 4D (right). Adjacency plot of CGs overlapping specified regions. Top panel is significance at individual CG level. Beta plot shows mean beta value for each group. Delta beta below shows mean beta value for each group relative to female values. Bottom plot shows in grey bars which layer a DMR was called in.

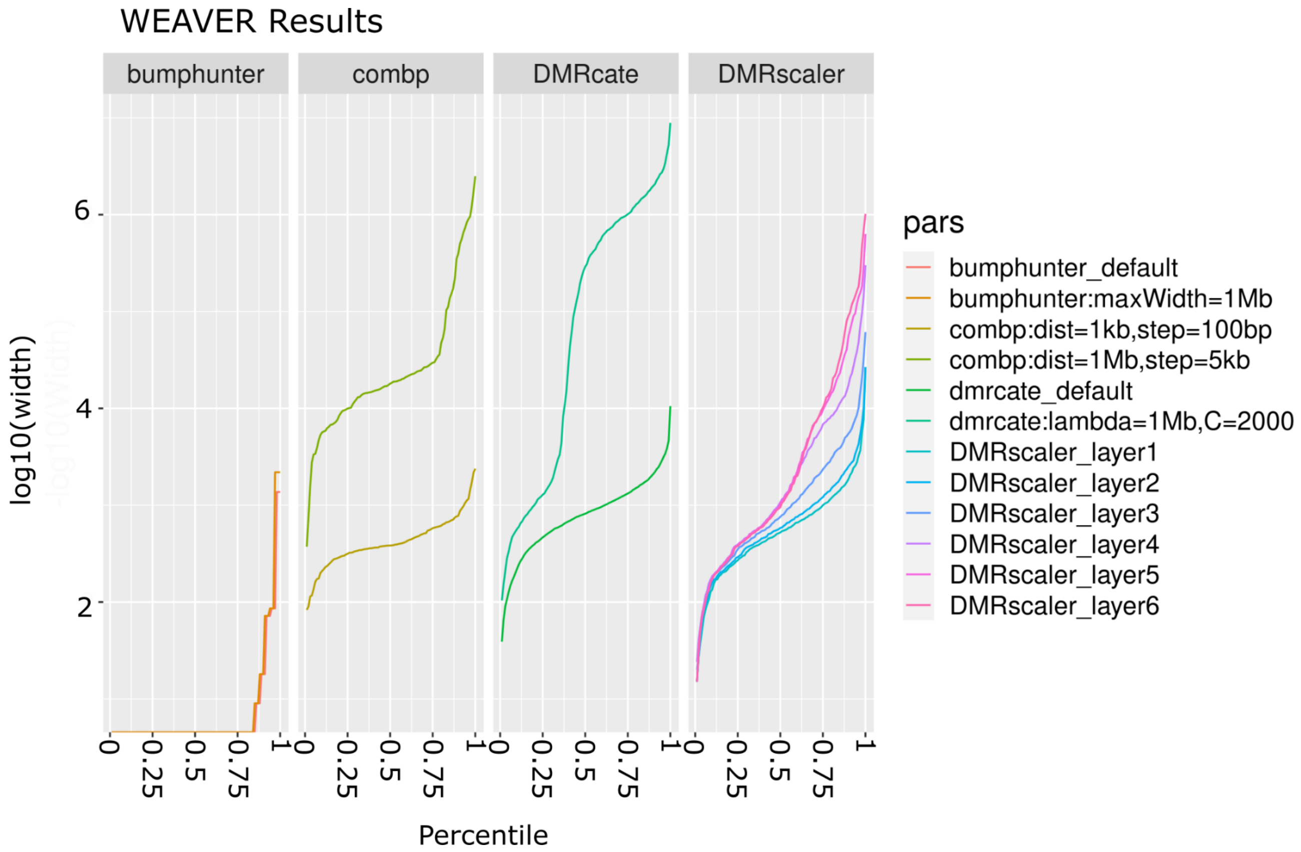


**FIGURE S14:** Weaver Analysis: Called DMR size percentiles

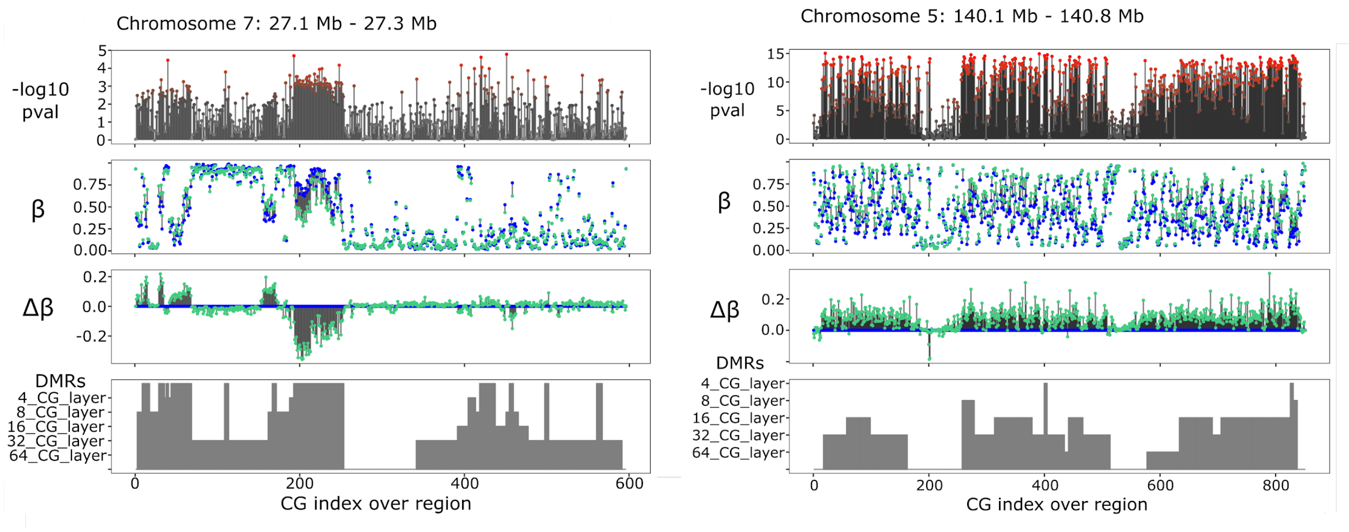


**Figure S15:** Weaver (left) and Sotos (right) analyses. Supplement to Figure 5B (left), 5D (right). Adjacency plot of CGs overlapping specified regions. Top panel is significance at individual CG level. Beta plot shows mean beta value for each group. Delta beta below shows mean beta value for each group relative to female values. Bottom plot shows in grey bars which layer a DMR was called in.

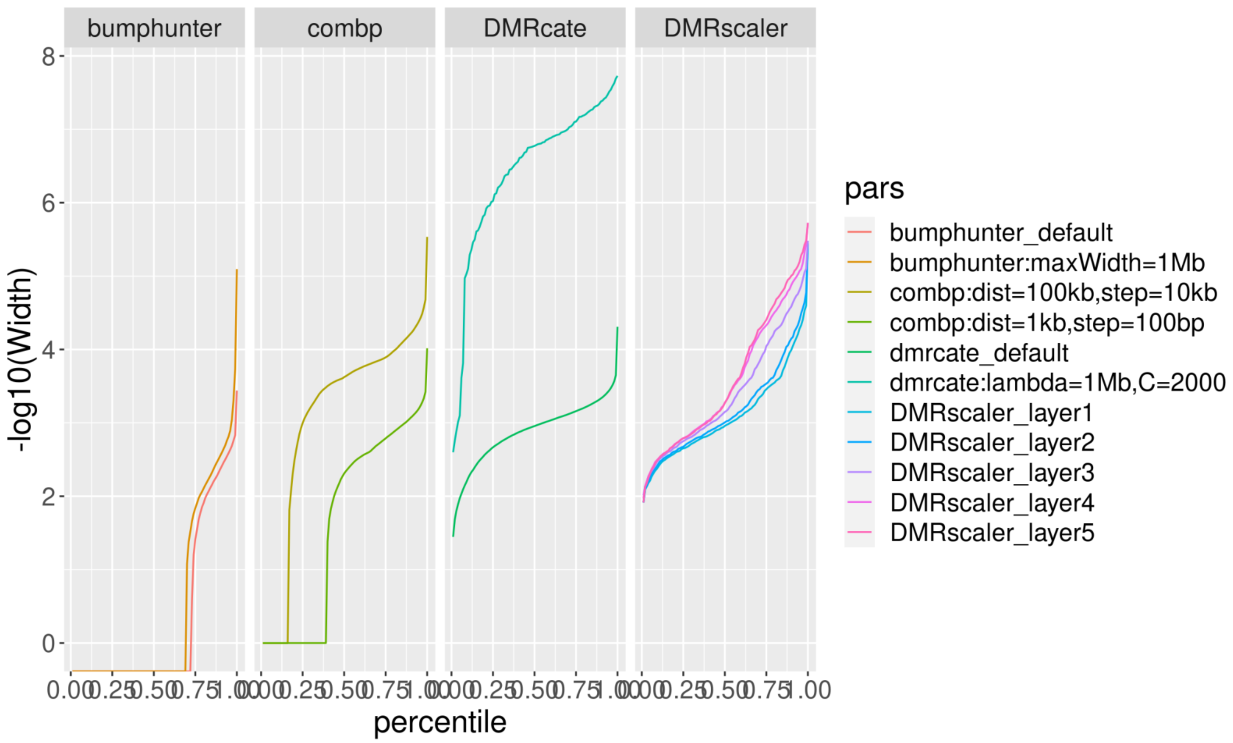


**FIGURE S16:** Sotos Analysis: Called DMR size percentiles.

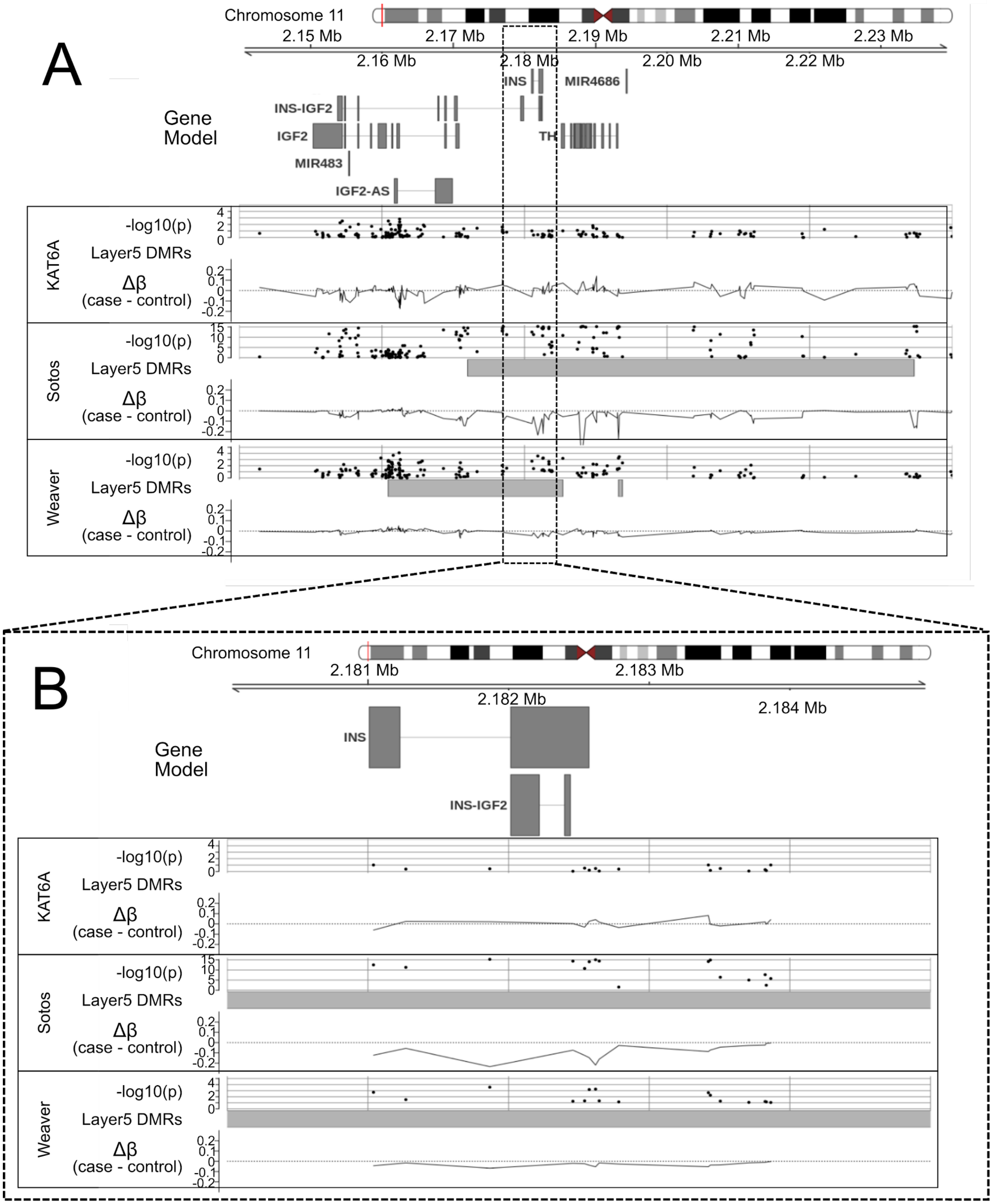


**Figure S17: *INS, IGF2, INS-IGF2* region.** Overlap between the Sotos and Weaver Syndromes, two overgrowth syndromes, identified a region proximal to and overlapping *INS, IGF2*, and

*INS-IGF2*.

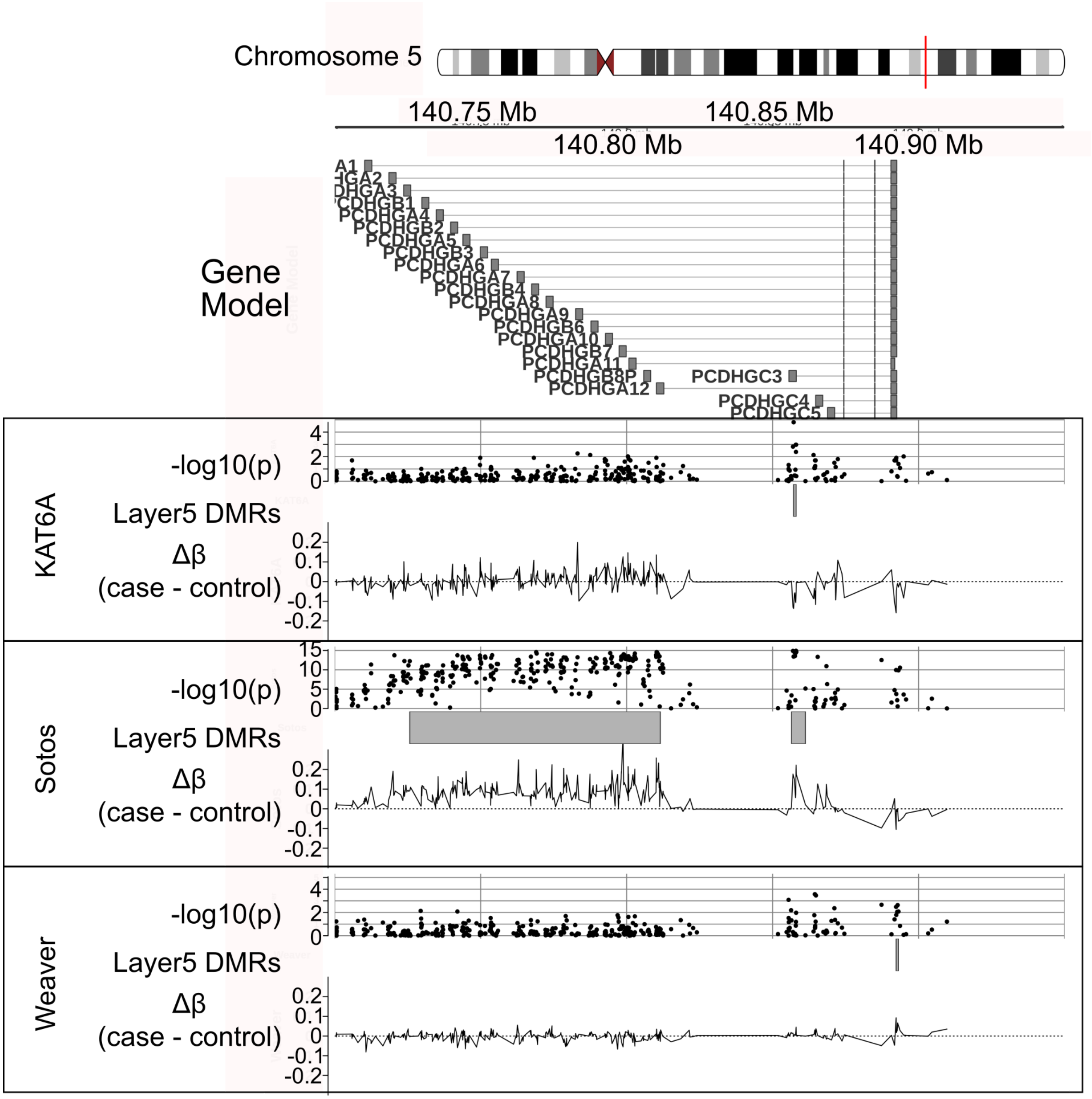


**Supplemental Figure 18 : *PCDHG* gene cluster GVIZ plot.** PCDHG genes were identified as genes overlapped by some DMR in each of the syndrome datasets analyzed. Within each box, top track is -log10(p) from wilcox test of beta values between cases and controls, middle track is DMRs called in the top level (layer5 aka 64 Adjacent CG layer) chosen as the top level is the most inclusive layer. Bottom track is the delta beta value (mean case Beta - mean control Beta).

| Analysis | pval cutoff (from permutation) | # CGs measured (Approximate) | Layer | Expected # sig windows | # DMRs Called | max FDR |
|---|---|---|---|---|---|---|
| Sex | 1/10,000 | 450,000 | 4 Adj CGs | 11.25 | 308 | 0.037 |
| KAT6A | 1/10,000 | 850,000 | 4 Adj CGs | 21.25 | 644 | 0.033 |
| WEAVER | 1/10,000 | 450,000 | 4 Adj CGs | 11.25 | 551 | 0.020 |
| SOTOS | 1/10,000 | 450,000 | 4 Adj CGs | 11.25 | 973 | 0.012 |

**Table S1:** Empiric choice of window selection significance level.

| KAT6A samples | genomic position | coding change (NM_006766.3) | protein change (NP_006757.2) |
|---|---|---|---|
| Patient 1 | chr8:g.41792353G>A | c.3385C>T | p.R1129* |
| Patient 2 | chr8:g.41795056G>A | c.3070C>T | p.R1024* |
| Patient 3 | chr8:g.41792353G>A | c.3385C>T | p.R1129* |
| Patient 4 | chr8:g.41791630C>A | c.4108G>T | p.E1370* |
| Patient 5 | chr8:g.41834753G>T | c.1136C>G | p.S379* |
| Patient 6 | chr8:g.41794839_41794840insG | c.3286_3287insC | p.C1096Sfs*6 |
| Patient 7 | chr8:g.41791376dupC | c.4362dupG | p.T1455Dfs*9 |
| Patient 8 | chr8:g.41791085A>C | c.4653T>G | p.S1551R |

**Table S2:** KAT6A Syndrome Patient Mutations

| **delta Beta = 0.1** | **AUCPR feature** | **AUCPR basepair** | **Simulated v Called Width Pearson's R** |
|---|---|---|---|
| bumphunter: maxGap=1e6 | 0 | 0 | -0.12 |
| comb-p: dist=1e6, step=5e3 | 0.34 | 0.68 | 0.66 |
| DMRcate: lambda=1e6,C=2e3 | 0.61 | 0.83 | 0.55 |
| DMRscaler | 0.86 | 0.97 | 0.95 |
| **delta Beta = 0.2** | | | |
| bumphunter: maxGap=1e6 | 0.14 | 0.09 | 0.00 |
| comb-p: dist=1e6, step=5e3 | 0.33 | 0.67 | 0.77 |
| DMRcate: lambda=1e6,C=2e3 | 0.79 | 0.90 | 0.67 |
| DMRscaler | 0.87 | 0.97 | 0.96 |
| **delta Beta = 0.4** | | | |
| bumphunter: maxGap=1e6 | 0.35 | 0.25 | 0.01 |
| comb-p: dist=1e6, step=5e3 | 0.35 | 0.70 | 0.81 |
| DMRcate: lambda=1e6,C=2e3 | 0.86 | 0.85 | 0.68 |
| DMRscaler | 0.88 | 0.97 | 0.98 |

**Table S3**: Results of variable value of delta Beta from simulation.

| Syndrome Pair | Layer1 CGs in DMR | Layer2 CGs in DMR | Layer3 CGs in DMR | Layer4 CGs in DMR | Layer5 CGs in DMR |
|---|---|---|---|---|---|
| **KAT6A** | 1194 | 1303 | 1700 | 2159 | 3035 |
| **Sotos** | 4957 | 5537 | 9338 | 11843 | 13141 |
| **Weaver** | 2162 | 2410 | 3974 | 6231 | 8711 |
| **KAT6A - Sotos** | 52 | 62 | 88 | 99 | 136 |
| **KAT6A - Weaver** | 22 | 32 | 38 | 121 | 246 |
| **Sotos - Weaver** | 208 | 228 | 370 | 414 | 453 |
| **KAT6A - Sotos - Weaver** | 0 | 0 | 0 | 0 | 0 |

**Table S9: Raw Count of Measured CGs in DMRs called by *DMRscaler* :** Layer1,2,3,4,5 are equivalent to 4,8,16,32,64 Adjacent CG Layers Respectively. CGs in DMR in each syndrome at each layer. Where multiple syndromes are listed, count represents CGs overlapped by some DMR in measured in each method using *DMRscaler*. Only the 425,733 Measured CGs present on both the Illumina 450k array, used for Sotos and Weaver, and the Illumina EPIC 850k array were used for overlap analysis.

| Syndrome Pair | Layer1 OR (OR 95% CI) | Layer2 OR (OR 95% CI) | Layer3 OR (OR 95% CI) | Layer4 OR (OR 95% CI) | Layer5 OR (OR 95% CI) |
|---|---|---|---|---|---|
| **KAT6A - Sotos** | 3.9 (2.95-5.15) | 3.82 (2.96-4.94) | 2.45 (1.97-3.04) | 1.69 (1.38-2.06) | 1.48 (1.24-1.76) |
| **KAT6A - Weaver** | 3.71 (2.42-5.66) | 4.47 (3.14-6.36) | 2.44 (1.77-3.37) | 4.06 (3.37-4.88) | 4.32 (3.78-4.93) |
| **Sotos - Weaver** | 9.39 (8.11-10.86) | 8.23 (7.16-9.45) | 4.73 (4.24-5.27) | 2.54 (2.3-2.81) | 1.75 (1.59-1.92) |

**Table S10: Odds Ratio (OR) for CGs found in DMR at each Layer of *DMRscaler* between all pairs of syndromes :** Layer1,2,3,4,5 are equivalent to 4,8,16,32,64 Adjacent CG Layers Respectively. Odds ratios (OR) are computed by labelling each measured CG as either in a DMR or not in a DMR for each syndrome to create a 2x2 contingency table to perform the odds ratio test on. Counts in Table S10. An confidence interval (CI) of the OR overlapping 1 implies no significant enrichment of CGs from one syndrome in the other.

| KAT6A Analysis DMR Summary Table | | | | |
|---|---|---|---|---|
| | Autosomes | | | |
| method | # DMRs | mean DMR width | median DMR width | % total width |
| **dmrscaler** | | | | |
| layer 1: 4 adjacent CpGs | 637 | 31.63 kb | 1.03 kb | 0.70% |
| layer 2: 8 adjacent CpGs | 599 | 57.61 kb | 1.09 kb | 1.2% |
| layer 3: 16 adjacent CpGs | 595 | 110.26 kb | 1.82 kb | 2.3% |
| layer 4: 32 adjacent CpGs | 575 | 173.70 kb | 2.19 kb | 3.5% |
| layer 5: 64 adjacent CpGs | 561 | 199.76 kb | 2.08 kb | 3.9% |
| **bumphunter** | | | | |
| default: maxGap = 500 bp | 3802 | 19 bp | 0 bp | 0.0025% |
| maxGap = 1 Mb | 3703 | 120 bp | 0 bp | 0.015% |
| **combp** | | | | |
| dist = 1kb, step = 100 bp | 72 | 359 bp | 334 bp | 0.00090% |
| dist = 1 Mb, step = 5 kb | 80 | 34.39 kb | 4.16 kb | 0.096% |
| **dmrcate** | | | | |
| default: lambda = 1 kb, C = 2 | 63 | 754 bp | 659 bp | 0.0016% |
| lambda = 1 Mb, C = 2000 | 51 | 34.78 kb | 673 bp | 0.062% |

**Table S11** : Summary of KAT6A analysis results

| WEAVER Analysis DMR Summary Table | | | | |
|---|---|---|---|---|
| | Autosomes | | | |
| method | # DMRs | mean DMR width | median DMR width | % total width |
| **dmrscaler** | | | | |
| layer 1: 4 adjacent CpGs | 547 | 1.16 kb | 582 bp | 0.022% |
| layer 2: 8 adjacent CpGs | 531 | 2.48 kb | 758 bp | 0.046% |
| layer 3: 16 adjacent CpGs | 518 | 9.05 kb | 1.07 kb | 0.16% |
| layer 4: 32 adjacent CpGs | 485 | 19.92 kb | 1.06 kb | 0.34% |
| layer 5: 64 adjacent CpGs | 472 | 33.12 kb | 956 bp | 0.54% |
| **bumphunter** | | | | |
| default: maxGap = 500 bp | 34 | 46 bp | 0 bp | 0.000054% |
| maxGap = 1 Mb | 33 | 72 bp | 0 bp | 0.000082% |
| **combp** | | | | |
| dist = 1kb, step = 100 bp | 220 | 487 bp | 384 bp | 0.0037% |
| dist = 1 Mb, step = 5 kb | 281 | 125.19 kb | 18.02 kb | 1.2% |
| **dmrcate** | | | | |
| default: lambda = 1 kb, C = 2 | 2932 | 1.06 kb | 817 bp | 0.11% |
| lambda = 1 Mb, C = 2000 | 837 | 721.93 kb | 290.22 kb | 21% |

**Table S12**: Summary of Weaver analysis results

| SOTOS Analysis DMR Summary Table | | | | |
|---|---|---|---|---|
| | Autosomes | | | |
| method | # DMRs | mean DMR width | median DMR width | % total width |
| **dmrscaler** | | | | |
| layer 1: 4 adjacent CpGs | 973 | 3.97 kb | 908 bp | 0.13% |
| layer 2: 8 adjacent CpGs | 976 | 5.47 kb | 1.01 kb | 0.19% |
| layer 3: 16 adjacent CpGs | 913 | 13.05 kb | 1.41 kb | 0.41% |
| layer 4: 32 adjacent CpGs | 854 | 24.09 kb | 1.88 kb | 0.71% |
| layer 5: 64 adjacent CpGs | 757 | 30.47 kb | 1.87 kb | 0.80% |
| **bumphunter** | | | | |
| default: maxGap = 500 bp | 6598 | 62 bp | 0 bp | 0.014% |
| maxGap = 1 Mb | 6276 | 312 bp | 0 bp | 0.068% |
| **combp** | | | | |
| dist = 1kb, step = 100 bp | 34033 | 417 bp | 204 bp | 0.49% |
| dist = 100kb, step = 10 kb | 20687 | 6.91 kb | 4.20 kb | 5.0% |
| **dmrcate** | | | | |
| default: lambda = 1 kb, C = 2 | 26701 | 1.09 kb | 903 bp | 1.0% |
| lambda = 1 Mb, C = 2000 | 255 | 9.13 Mb | 5.94 Mb | 81% |

**Table S13** : Summary of Sotos analysis results
